# A Protein Blueprint of the Diatom CO_2_-Fixing Organelle

**DOI:** 10.1101/2023.10.26.564148

**Authors:** Onyou Nam, Sabina Musial, Manon Demulder, Caroline McKenzie, Adam Dowle, Matthew Dowson, James Barrett, James N. Blaza, Benjamin D. Engel, Luke C. M. Mackinder

## Abstract

Diatoms are central to the global carbon cycle. At the heart of diatom carbon fixation is an overlooked organelle called the pyrenoid, where concentrated CO_2_ is delivered to densely packed Rubisco. Diatom pyrenoids fix approximately one-fifth of global CO_2_, but virtually nothing is known about this organelle in diatoms. Using large-scale fluorescence protein tagging and affinity purification-mass spectrometry, we generate a high-confidence spatially-defined protein-protein interaction network for the diatom pyrenoid. Within our pyrenoid interaction network are 10 proteins with no known function. We show that six of these form a static shell encapsulating the Rubisco matrix of the pyrenoid, with the shell critical for pyrenoid structural integrity, shape, and function. Although not conserved at a sequence level, the diatom pyrenoid shares some architectural similarities to prokaryotic carboxysomes. Collectively, our results support the convergent evolution of pyrenoids across the two main plastid lineages and uncover a major structural and functional component of global CO_2_ fixation.

## Introduction

Approximately one-third of global carbon fixation takes place in pyrenoids.^1^ Pyrenoids are biomolecular condensates of the principal CO_2_-fixing enzyme Rubisco found in the chloroplasts of algae.^2^ There are two major chloroplast lineages, the green and red plastids with algae within these lineages proposed to have convergently evolved pyrenoids.^3,4^ Nearly all of our knowledge of pyrenoid function and composition comes from the model terrestrial green plastid alga *Chlamydomonas reinhardtii.*^2,5^ *C. reinhardtii* is a powerful model organism^6^ but global carbon fixation is primarily driven by oceanic red plastid containing algae, such as diatoms, where our knowledge is still in its infancy.^7–9^ Diatoms are responsible for up to 20% of global net primary production in the modern ocean, are estimated to fix ∼70 gigatons CO_2_ yr^-1^, and are fundamental for the long-term storage of carbon by driving the flux of organic material from the ocean surface to sediments.^10,11^

Pyrenoids are found at the heart of algal CO_2_ concentrating mechanisms (CCMs).^2,7^ CCMs overcome the slow diffusion of CO_2_ in water and the catalytic limitations of Rubisco by actively pumping inorganic carbon from the external environment into the cell and releasing it as CO_2_ in the pyrenoid where it can be fixed by tightly packaged Rubisco. In *C. reinhardtii* a disordered linker protein, EPYC1, condenses Rubisco to form the liquid-liquid phase separated matrix of the pyrenoid.^1,12,13^ A shared Rubisco binding motif found in EPYC1 and numerous other pyrenoid components enables targeting to and structural organization of the pyrenoid. With proteins containing this motif proposed to link the matrix to specialized traversing thylakoids called pyrenoid tubules and link the matrix to the surrounding starch sheath.^14^ Inorganic carbon in the form of HCO_3_^-^ is shuttled into the pyrenoid tubules by bestrophin-like proteins^15^ where a carbonic anhydrase converts it to membrane permeable CO_2_ that can leak out and be fixed by Rubisco in the pyrenoid matrix.^5,15,16^ The surrounding starch acts as a diffusion barrier to minimize CO_2_ leakage out of the pyrenoid.^17,18^ Fluorescent protein tagging,^14,19^ affinity purification followed by mass spectrometry (APMS)^19^ and proximity labeling^20^ have enabled a high-confidence pyrenoid proteome to be determined with multiple components now functionally characterized. This has enabled *C. reinhardtii* proto-pyrenoid engineering in plants^21^ and a parts-list of components that should theoretically enable the engineering of a functional pyrenoid-based CCM to enhance plant photosynthesis.^17,22^

Diatom pyrenoids have some shared features with the *C. reinhardtii* pyrenoid including condensed Rubisco and traversing thylakoids, it is also proposed that they may function analogously with some, although limited, conservation of inorganic carbon delivery proteins.^7,8^ However, it is still unclear the level of conservation of structural proteins.^9^ The only structural component identified so far for the diatom pyrenoid is PYCO1 a Rubisco linker protein found in the pyrenoid of the pennate diatom *Phaeodactylum tricornutum* and suggested to be responsible for phase separating Rubisco to form the pyrenoid matrix.^22,23^ However, PYCO1 is not widely conserved, being absent in centric diatoms, and its functional importance is yet to be determined. Outside of PYCO1 most previous diatom CCM research has focused on inorganic carbon uptake. In *P. tricornutum* several candidates belonging to the SLC4 family of transporters have been proposed for HCO_3_^-^ uptake at the plasma and chloroplast membranes.^23,24^ Bestrophin-like proteins are also implicated in the shuttling of HCO_3_^-^ into the thylakoid lumen where a θ-type carbonic anhydrase has been identified that is thought to function by releasing CO_2_ from the thylakoid membranes that traverse the pyrenoid matrix.^25,26^ Less is known about inorganic carbon uptake in the centric diatom *Thalassiosira pseudonana*. SLC4 candidates have been implicated in inorganic carbon uptake across both the plasma and thylakoid membranes.^8,27^ Recently two bestrophin-like proteins were localized to the *T. pseudonana* pyrenoid, likely in the pyrenoid penetrating thylakoid (PPT)^28,29^ and a θ-type carbonic anhydrase 2 confirmed to be located in the PPT.^18^ Although, supporting functional data is missing for these proteins in *T. pseudonana*. Absent from diatom pyrenoids is a starch sheath encapsulating the pyrenoid. In *C. reinhardtii* flux balance modeling of pyrenoid function^17^ and analysis of starch mutants^28^ indicate that a diffusion barrier is essential for efficient pyrenoid function to minimize CO_2_ leakage. How the diatom pyrenoid minimizes CO_2_ leakage is a substantial outstanding question.

Here we rapidly advance our knowledge of the diatom pyrenoid by developing an iterative fluorescent protein tagging followed by APMS approach in *T. pseudonana*, which belongs to a global biogeochemically important genus. These data enabled us to build a high-confidence diatom pyrenoid interaction network, identifying multiple new pyrenoid proteins, many with no previously known functional domains. A family of these proteins form a static shell that encapsulates the Rubisco matrix and is critical for pyrenoid shape, structural integrity, and CCM function. Our findings provide new insight into a globally important organelle and provide additional molecular parts for engineering a CCM into crop plants to improve productivity.

## Results and discussion

### Rubisco co-immunoprecipitation mass spectrometry to identify diatom pyrenoid components

Although diatoms play a central role in global biogeochemical cycles, very little is known about the diatom pyrenoid. In *T. pseudonana* cells have two chloroplasts, each surrounded by four membranes, with each chloroplast containing a single centrally positioned lenticular-shaped pyrenoid (Fig. 1A). As a starting point to understand *T. pseudonana* pyrenoid composition, we performed co-immunoprecipitation coupled with mass spectrometry (coIPMS), using the main pyrenoid component, Rubisco, as a bait protein (Fig. 1B). To immunoprecipitate Rubisco, we used an antibody raised to a conserved 12 amino acid surface-exposed peptide on the Rubisco large subunit (rbcL) (Fig. S1). By comparing two independent coIPMS experiments, consisting of two and three technical replicates, respectively, against non-antibody control experiments, we identified 36 putative pyrenoid components out of a total of 1167 detected proteins identified with two or more spectral counts (Table S1). For these exploratory experiments, we applied a relaxed cut-off based on the fold-change enrichment of bestrophin-like protein 2 (BST2) that we had previously localized to the pyrenoid^28^ and would expect to only have a weak enrichment due to it being predicted to be a membrane protein (Fig. 1B). Top hits from our rbcL coIPMS experiments were then fed into an iterative fluorescent protein tagging, localization, and APMS framework that we used to rapidly build a spatially defined pyrenoid proteome (Fig. 1C).

**Figure 1.**
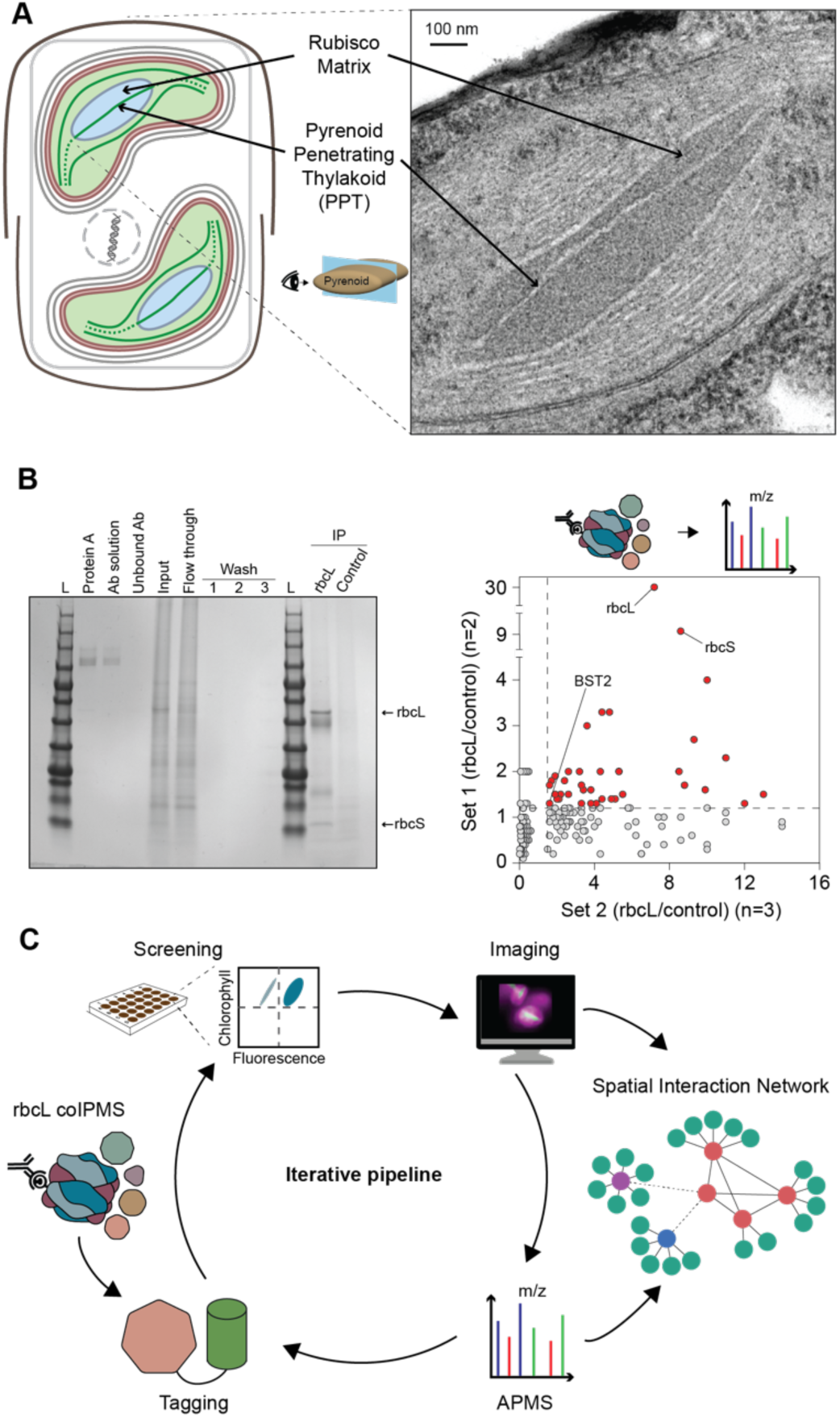
Identification of candidate diatom pyrenoid components. A. The diatom *T. pseudonana* has two chloroplasts each containing a single pyrenoid traversed by a specialized membrane called the pyrenoid penetrating thylakoid (PPT). The connection between the PPT and the wider thylakoid network is unresolved. B. Rubisco co-immunoprecipitation followed by mass spectrometry (coIPMS) to identify candidate pyrenoid components. Left: example Rubisco large subunit (rbcL) coIP with different steps resolved by SDS-PAGE. Right: Two experiments consisting of either two or three technical coIPs were performed and the antibody bound fractions submitted for mass spectrometry. Cut-offs (dashed lines) were determined based on bestrophin-like protein 2 (BST2) that was previously shown to be pyrenoid localized. Ab, antibody; L, protein ladder. C. An iterative pipeline to determine the pyrenoid spatial interaction network. RbcL coIP data fed into a high-throughput fluorescence protein tagging pipeline. Pyrenoid confirmed proteins were then used for affinity purification followed by mass spectrometry (APMS). Interactors were tagged to confirm pyrenoid localization and used in subsequent APMS rounds.

### Development of a high-throughput tagging pipeline in *T. pseudonana* identifies multiple novel pyrenoid components

To enable rapid cycling through our iterative pipeline, we set out to establish high-throughput fluorescent protein tagging and screening in diatoms. We initially adapted our Golden Gate Modular Cloning-based episomal assembly framework^30^ to be 96-well compatible and combined it with multi-well diatom transformation via bacterial conjugation. We coupled this with 48-well plate strain maintenance and 96-well plate flow cytometry screening for clonal fluorophore-fusion expressing lines (Fig. 2A). As coIPMS data on a small-scale is inherently noisy with both false positives and false negatives^31^ we applied our tagging pipeline to 22 Rubisco coIPMS hits to validate if they were bona fide pyrenoid proteins. Nourseothricin-positive transformants were picked and screened for positive fluorescence using flow cytometry. Due to typical mosaic colony presence^12^, either multiple rounds of screening or screening of several independent colonies was required to identify stable mEGFP expressing lines. We found that screening 8–12 colonies would typically yield a stable cell population with >90% of cells mEGFP positive (Fig. S2). Positive lines were subsequently imaged by confocal microscopy (Fig. 2B, Fig. S3 and S4). From the initially identified Rubisco-interacting proteins, we saw 13 proteins localizing to distinct sub-regions of the pyrenoid (Fig. 2B and Fig. S4; see below for further discussion). However, 9 candidates were either localized to chloroplast sub-regions adjacent to the pyrenoid (4), or to distinct non-chloroplast regions (5) (Fig. S3) indicating that our coIPMS data contains false positives.

**Figure 2.**
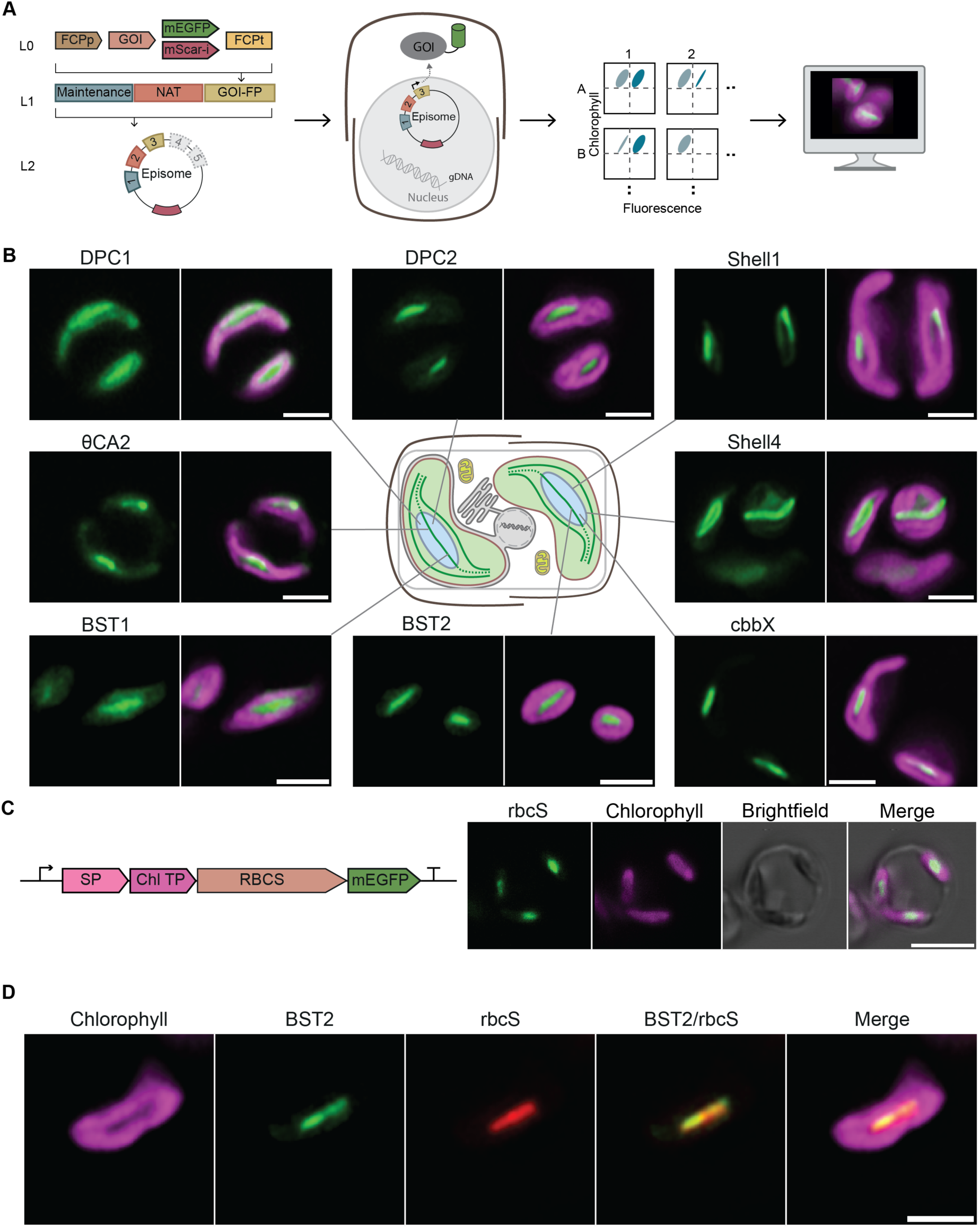
High-throughput fluorescence protein tagging identifies multiple new diatom pyrenoid components. A. Candidate genes were rapidly cloned using Golden Gate modular cloning, transformed into *T. pseudonana* via bacterial conjugation and screened by flow cytometry for fluorescence prior to imaging. Gene of interest (GOI) expression was driven by the fucoxanthin chlorophyll a/c-binding protein promoter (FCPp) and terminated using the FCP terminator (FCPt). B. Multiple proteins localized to the canonical pyrenoid region within the chloroplast. Newly identified pyrenoid components with no sequence-predicted function are termed Diatom Pyrenoid Components (DPCs) or Shells. Green: mEGFP fusion protein; Magenta: chlorophyll autofluorescence. Scales bars: 2 µm. C. Development of a pyrenoid matrix marker. By using the BST2 signal and transit peptide sequences we could express chloroplast encoded rbcS from nuclear episomes and target it to the chloroplast. Scale bar: 5µm. D. Co-localization of BST2-mEGFP (green) and rbcS-mScarlet-I (red) expressed from the same episome. Scale bar: 2 µm.

To accurately determine sub-pyrenoid localization of components, we developed a dual-tagging approach using two spectrally compatible fluorophores. We first developed a pyrenoid matrix marker for co-localization. In green algae, nuclear-encoded rbcS-fluorescent protein fusions have been powerful for understanding sub-pyrenoid spatial organization^31^ and for determining the liquid-like properties of the pyrenoid.^32^ As the rbcS of diatoms is chloroplast encoded, and no *T. pseudonana* chloroplast transformation protocol is available, we wondered if we could target an episomal expressed rbcS-mEGFP to the pyrenoid. Using the N-terminal signal and transit peptide sequences from the nuclear encoded chloroplast localized BST2 protein,^28^ we successfully targeted rbcS to the pyrenoid, allowing us to clearly define the pyrenoid matrix (Fig. 2C). Second, we tested assembling two target genes with different fluorophores on the same episome. We decided to initially validate the BST2 localization by making an episome with BST2-mEGFP and rbcS-mScarlet-I, a fluorophore we had previously validated using our system.^33^ Using this approach, we observed a clear pyrenoid localization of BST2, with the rbcS signal extending outside that of BST2, supporting a PPT localization of BST2 (Fig. 2D).

### Establishing large-scale APMS to build a pyrenoid interaction network

To rapidly expand the pyrenoid proteome we developed and optimized an APMS pipeline in *T. pseudonana* and used our GFP-tagged pyrenoid proteins as baits in this pipeline. Lines expressing GFP-tagged proteins were typically grown at ambient CO_2_, where the CCM is fully active.^19,34^ In triplicate, GFP-trap nanobodies were used to enrich for target proteins from cell lysate and their interactors determined via LC-MS/MS (Fig. 3A, Table S2). Protein-protein interactions were stringently defined by comparing both CompPASS^35^ and SAINT analysis^28^ scores (Fig. 3A, Table S3) that use different weighting criteria to identify true interactors from non-specific background using label-free proteomic quantitation data. We set interaction confidence thresholds based on the known interaction of rbcS with rbcL, which resulted in proteins in the top 2.2% for CompPASS and top 1% for SAINT being designated as high confidence interactors.

**Figure 3.**
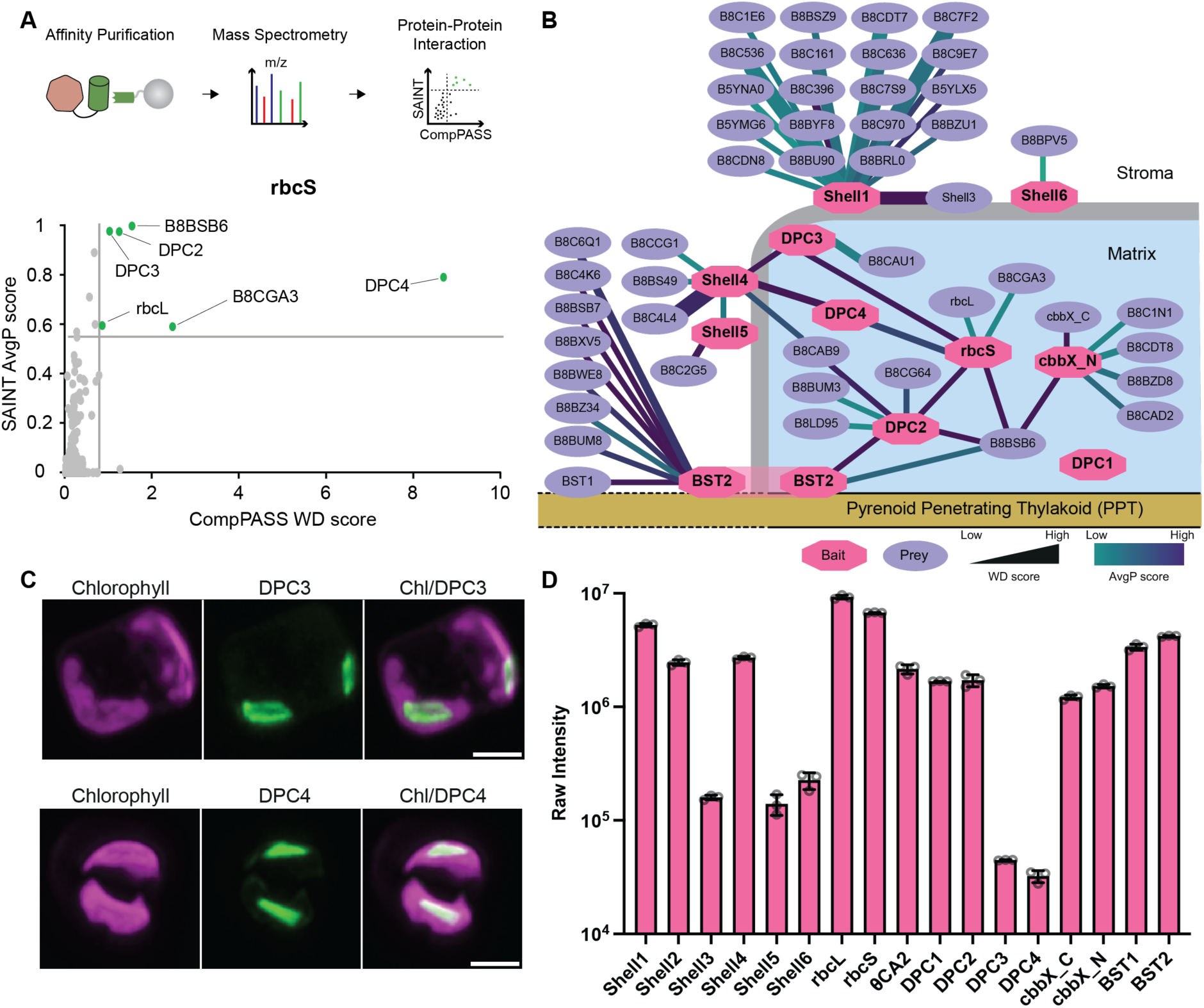
A spatial interaction network of the diatom pyrenoid. A. Top: mEGFP tagged proteins were affinity purified using GFP nanobodies, interactors identified using mass spectrometry and high-confidence protein-protein interactions determined using SAINT and CompPASS analysis. Bottom: analysis example for RbcS APMS. RbcS is not present in the plot due to bait spectral counts being set to zero prior to SAINT and CompPASS analysis. B. Spatially determined protein-protein interaction network of the pyrenoid. Where available, positioning is based on localization determined by confocal microscopy. Due to 96% similarity of the mature proteins of Shell1 and Shell2 all peptides were assigned to Shell1 in the APMS study. C. Confocal localization of DPC3 and DPC4 confirming pyrenoid localization (max intensity z-stack projections). Green: mEGFP fusion protein; Magenta: chlorophyll. Scale bars: 2µm. D. Whole cell mass spectrometry raw intensity values of pyrenoid confirmed proteins. Data is from wildtype cells ran in triplicate. Error bars: S.D. of the mean.

Our initial 13 pyrenoid localized proteins included the Rubisco small subunit rbcS; BST1 and BST2 that are bestrophin-like proteins proposed to be involved in HCO_3_^-^ uptake into the PPTs;^25,26^ θCA2 most likely involved in CO_2_ release from HCO_3_^-^ within the PPTs;^36^ cbbX a nuclear encoded red-type Rubisco activase^37^ that until now has not been localized in algae with red plastids;^1,38^ and 8 uncharacterized proteins with no clear function. These newly identified GFP-tagged pyrenoid components were called Diatom Pyrenoid Components 1 and 2 (DPC1 and DPC2) and Shell 1-6. Whilst DPC1 and DPC2 were predominantly in the pyrenoid matrix, the initial tagging of Shell1 and Shell4 showed that they may encapsulate the pyrenoid (Fig. 2B; see below). A subset of these pyrenoid-localized components were utilized for APMS using our iterative pipeline (Fig. 1C, Fig. 3B and Table S4). Subsequently, two additional components, DPC3 and DPC4, which had strong interactions with Shell4 and rbcS but again with no sequence predictable function, were localized and fed into our APMS pipeline. Whereas DPC4 was found throughout the pyrenoid with a matrix-type localization, DPC3 appeared to encapsulate the pyrenoid similar to the Shell proteins (Fig. 3C). Combining the data and using our stringent interaction scoring approach enabled us to expand and build a high-confidence pyrenoid interaction network for *T. pseudonana* built from 11 baits and containing 46 additional protein nodes linked by 57 interaction edges (Fig. 3B). In the network, interaction confidence can be further interpreted by CompPASS and SAINT score magnitude (line thickness and color respectively in Fig. 3B) as well as the number of connecting edges with baits.

RbcS and DPC2 appear to be key hub proteins, each linking four nodes that further link to confirmed pyrenoid components. How Rubisco is packaged into the pyrenoid is unknown in *T. pseudonana*. With the absence of an EPYC1 or PYCO1 homolog to phase separate Rubisco, it was hypothesized that an alternative repeat protein could be fulfilling this role. Unexpectedly, none of the pyrenoid proteins identified in our study contain a repeated sequence with the expected frequency of ∼60 amino acids,^39–41^ potentially indicating that pyrenoid assembly in *T. pseudonana* could be based on different biophysical principles to both green algal and pennate diatom pyrenoids. The localization of DPC2 solely to the pyrenoid (Fig. 2B), its Rubisco interaction (Fig. 3B), and its relatively high abundance from whole cell mass spectrometry (Fig. 3D and Table S5) make DPC2 a strong candidate for future studies.

### *T. pseudonana* has six Shell homologs, and Shell proteins are found across algal lineages containing red plastid secondary endosymbionts

In our initial tagging, we were intrigued to see that two proteins with no annotated functional domains appeared to encapsulate the pyrenoid matrix (Fig. 2B: Shell1, Shell4). In the well-characterized green algal pyrenoid, chloroplast synthesized starch forms a sheath that encapsulates the Rubisco matrix and acts as a CO_2_ leakage barrier to enhance CCM efficiency.^17,18^ In diatoms, the main carbohydrate storage molecule is chrysolaminaran, which is stored in cytosolic vacuoles^40–43^ with no clear carbohydrate barrier surrounding the diatom pyrenoid. Pyrenoid modeling of inorganic carbon fluxes has shown that a CO_2_ diffusion barrier is critical for an efficient CCM,^17^ thus opening up the question of how diatoms and other algal lineages that lack starch encapsulated pyrenoids operate efficient CCMs.^43–45^ Although protein encapsulation of pyrenoids has not previously been identified, cyanobacterial Rubisco-containing carboxysomes have a well-characterized protein shell that is proposed to act as a CO_2_ barrier to minimize leakage,^44–46^ and is essential for carboxysome biogenesis and shape.^46^ Instead of starch, we hypothesized a protein shell could have an analogous function in diatoms. We set out to further understand the importance of the Shell proteins for pyrenoid function.

BLAST analysis of Shell1 and Shell4 identified four additional homologs in the *T. pseudonana* genome that all contain two predicted beta-sheet domains (Fig. 4A and Fig. S5) and are predicted to be structurally similar to each other but distinct from DPC3 (Fig. S6). We explored the distribution of these proteins across different evolutionary lineages. BLAST analysis against the NCBI database identified homologs in stramenopiles (including other diatoms), pelagophytes and haptophytes; all of which are photosynthetic algae that contain red plastids originated from secondary endosymbiotic events (Fig. 4B and Table S6). However, Shell proteins are absent in the rhodophytes, the primary endosymbiotic red plastid lineage that was engulfed by a heterotrophic host to form the red plastid secondary endosymbiotic lineages. This suggests that Shell proteins were either not present in the engulfed red alga but may have been present in the genome of the heterotrophic host prior to endosymbiosis, were present in the engulfed red alga but have since been lost in rhodophytes, or evolved after engulfment. Given that Shells are not found in extant non-photosynthetic organisms and are not present in pyrenoid-containing rhodophytes, it seems plausible that they evolved following the secondary endosymbiotic event. Further supporting a role in pyrenoid function of Shell proteins, TEM data from available literature indicates that algae found within the Shell protein-containing clades all possess pyrenoids.^47^

**Figure 4.**
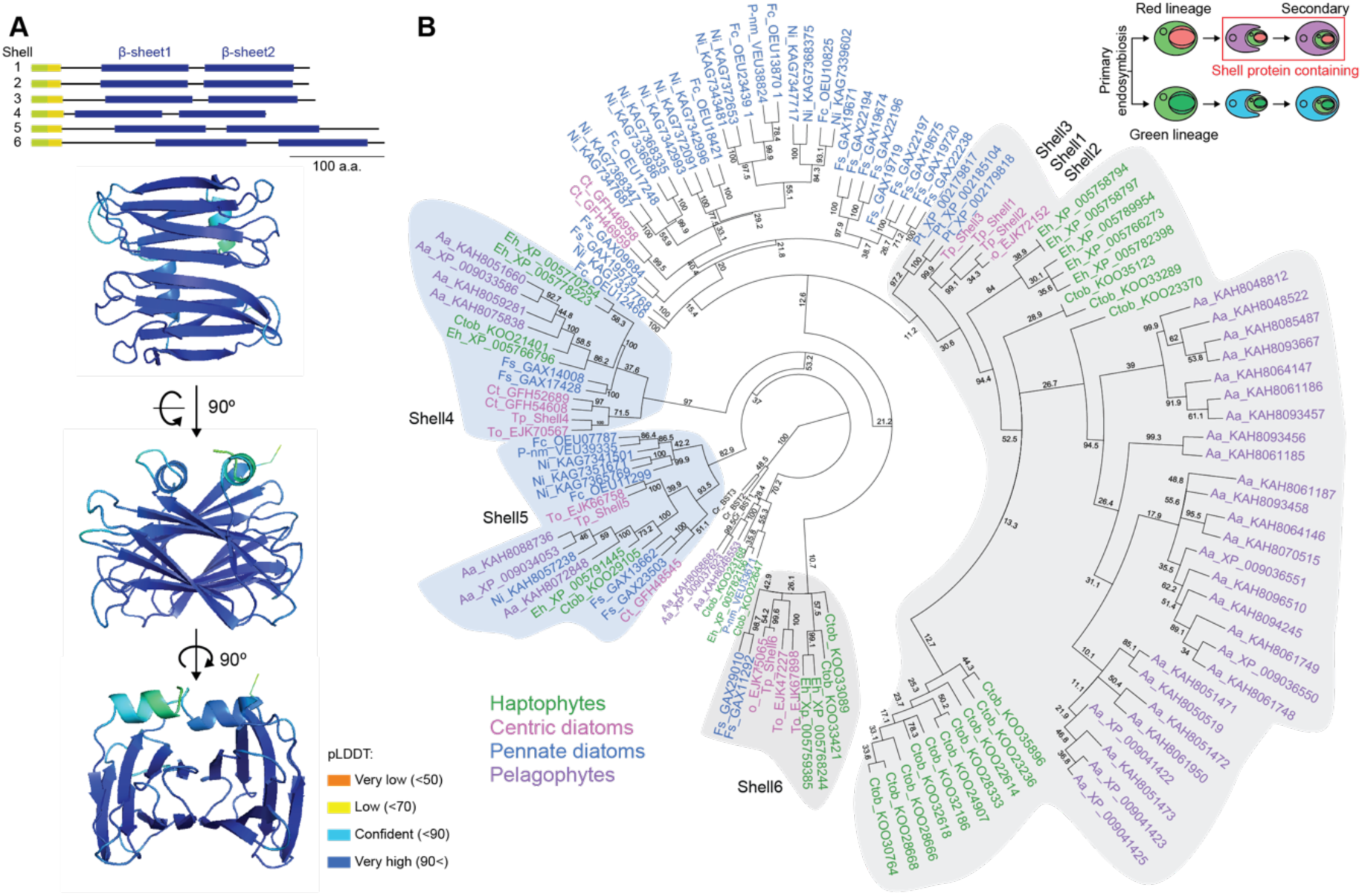
*T. pseudonana* has six Shell homologs and Shell proteins are found across red plastid secondary endosymbionts. A. *T. pseudonana* has six Shell protein homologs that all contain two beta-sheet regions. Top: domain alignment of Shell homologs, Green is signal peptide, yellow is transit peptide. Bottom: AlphaFold2 structure prediction of Shell1. pLDDT: predicted local distance difference test. The N-terminal and C-terminal disordered regions are not shown due to low confidence structural prediction. B. Phylogenetic analysis indicates that Shell proteins are found in red plastid secondary endosymbionts including stramenopiles, pelagophytes and haptophytes. Bootstrap values are shown. Cartoon inset represents the two main plastid lineages and where Shell proteins are present. Aa, *Aureococcus anophagefferens*; Ct, *Chaetoceros tenuissimus*; Ctob, *Chrysochromulina tobinii*; Eh, *Emiliania huxleyi*; Fs, *Fistulifera solaris*; Fc, *Fragilariopsis cylindrus*; Ni, *Nitzschia inconspicua*; Pt, *Phaeodactylum tricornutum*; P-nm, *Pseudo-nitzschia multistriata*; To, *Thalassiosira oceanica*; Tp, *Thlassiosira pseudonana*. *Chlamydomonas reinhardtii* bestrophins (Cr_BST1-3) have been used as an outgroup.

### Shell proteins encapsulate the pyrenoid matrix and localize to distinct sub-regions

To elucidate the precise sub-regions of the six Shell proteins, we co-expressed rbcS-mScarlet-I with mEGFP-tagged Shell proteins (Fig. 5A and Fig. S7). Analysis of fluorescence intensity of pyrenoid transects (Fig. 5B and Fig. S7) and max intensity z-stack projections (Fig. 5C and Fig. S7) indicate that all six proteins encapsulate the Rubisco matrix of the pyrenoid. To prioritize Shell proteins for further investigation, we looked at their relative abundance in our whole cell mass spectrometry data performed on wildtype cells (Fig. 3D and Table S5). The most abundant homologs were Shell1 and Shell2, which share 92% identity, and Shell4, which shares 34% identity with Shell1 (Fig. S5B). As the diatom pyrenoid shape can be generalized as an elliptic cylinder with curved ends (Fig. 1), the apparent localization of the Shell proteins is highly dependent on the orientation of the chloroplast and pyrenoid during imaging (Fig. 5A-C and Fig. S7), making the assignment to sub-pyrenoid surface regions challenging. To explore if Shell1 and Shell4 occupy the same regions of the shell we co-expressed Shell1 and Shell4 with different fluorescent tags. Imaging shows that they localize to distinct regions of the pyrenoid periphery, suggesting that they may have different roles in the formation or function of the shell (Fig. 5D-F).

**Figure 5.**
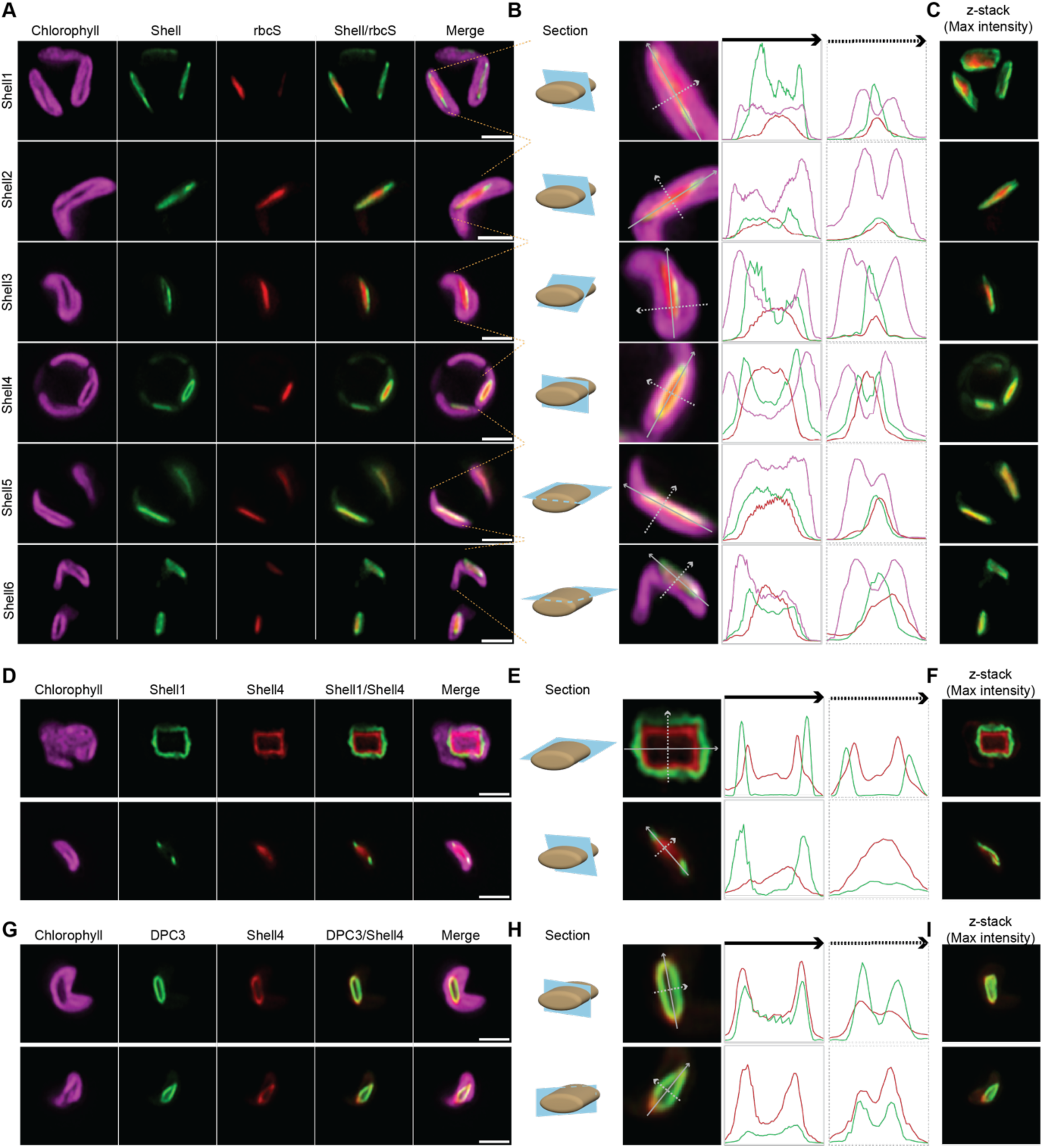
Shell proteins encapsulate the pyrenoid matrix, and Shell1 and Shell4 occupy distinct regions of the pyrenoid surface. A. Co-localization of Shell proteins with Rubisco. B. Fluorescence intensity cross-sections of chlorophyll, Shell and rbcS across the pyrenoid. C. Z-stack max intensity projections of C. D. Co-localization of Shell1 and Shell4 in two cells with different pyrenoid orientations. E. Fluorescence intensity cross-sections of Shell1 and Shell4 across the pyrenoid. F. Z-stack max intensity projections of E. G. Co-localization of DPC3 and Shell4 in two cells with different pyrenoid orientations. H. Fluorescence intensity cross-sections of DPC3 and Shell4 across the pyrenoid. I. Z-stack max intensity projections of H. Scale bars: 2µm. For B, E and H the level of Zoom changes between images.

Although Shell proteins were identified in our rbcL coIPMS and rbcS APMS data, they did not fall above the stringent thresholds we defined for high-confidence interactors with either rbcS or rbcL. Their pyrenoid-peripheral localization and low confidence Rubisco interaction suggest Shell proteins could interact indirectly with the Rubisco matrix. The shell-like pattern displayed by DPC3 (Fig. 3C) raises the possibility that this protein could be acting as an intermediary Rubisco matrix-shell adaptor or potentially an additional shell component. The co-expression of DPC3 with Shell4 shows that they co-localize (Fig. 5G-I). The relatively low protein abundance of DPC3 in comparison to Shell1 and Shell4 (Fig. 3D) indicates that DPC3 is most likely a minor protein constituent of the pyrenoid surface.

### The pyrenoid matrix and shell proteins have minimal mobility in the pyrenoid

To understand the dynamics of pyrenoid components, we leveraged our tagged lines. If the shell proteins form a lattice around the Rubisco matrix, similar to carboxysome shell proteins in assembled carboxsomes,^47,48^ then we would expect them to form a static layer. We investigated the mobility of Shell1 and Shell4 by fluorescence recovery after photobleaching (FRAP). Both Shell1 and Shell4 showed minimal rearrangement after photobleaching (Fig. 6A, B). Although this does not provide direct evidence that they form a CO_2_ diffusion barrier, the static nature of the pyrenoid shell is consistent with our current understanding of both cyanobacterial carboxysome shell and green algal starch sheath CO_2_ diffusion barriers.

**Figure 6:**
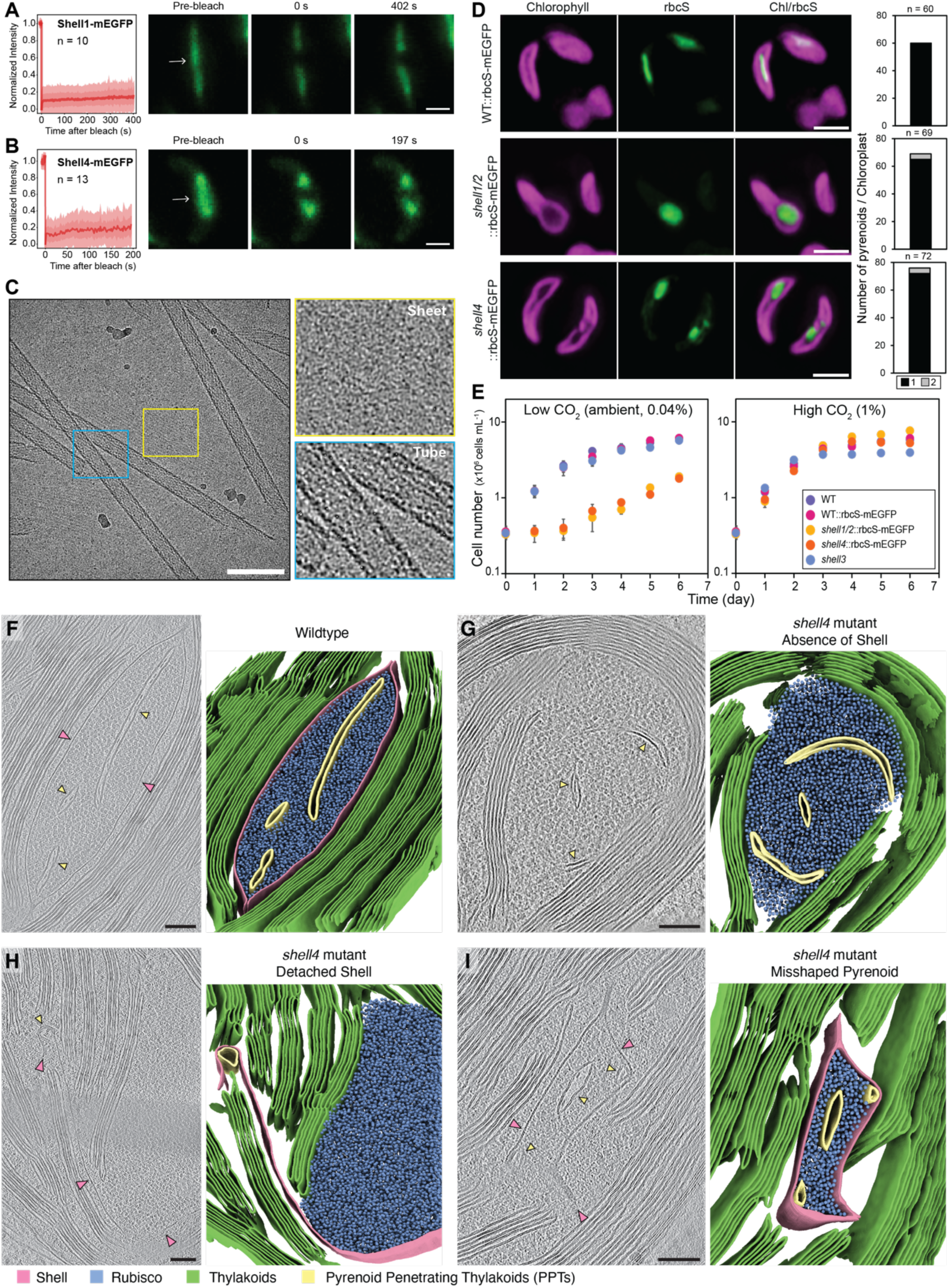
Shell proteins are static on the pyrenoid surface, can self-assemble into sheets and tubes, and are essential for CCM function and pyrenoid architecture. A-B. Fluorescence recovery after photobleaching (FRAP) experiments for Shell1 (A) and Shell4 (B). Arrows indicate photobleached regions. Scale bars: 1 µm. Plots show S.E.M. and S.D. of the mean. C. Shell1 can form tubes and sheets as seen by cryo-EM. Scale bar: 100 nm. D: Tagged rbcS in wildtype (WT::rbcS-mEGFP, top), a *shell1/2* mutant (*shell1/2*::rbcS-mEGFP, middle) and a *shell4* mutant line (*shell4*::rbcS-mEGFP, bottom). Scale bars: 2 µm. Bar charts indicate pyrenoid number per chloroplast in corresponding line. E: Growth rate of WT, WT::rbcS-mEGFP, *shell1/2*::rbcS-mEGFP, *shell4*::rbcS-mEGFP and a *shell3* mutant. F-I: Cryo-electron tomography of wildtype *T. pseudonana* (F) and the *shell4*::rbcS-mEGFP line (G-I). In each panel, the left image shows a 2D slice through the tomogram, and the right image shows the corresponding 3D segmentation of the Shell structure (pink), Rubisco matrix (blue), thylakoids (green), and PPTs (yellow). Scale bars: 100 nm.

In *C. reinhardtii*, the pyrenoid matrix has liquid-like properties. Rubisco, EPYC1 and Rubisco activase (RCA1) all show rapid rearrangement in FRAP experiments on the timescale of ∼30 s after photobleaching.^48,49^ In *T. pseudonana* Rubisco is densely packed in the pyrenoid but does not formed ordered arrays.^50^ Surprisingly, FRAP experiments on both *T. pseudonana* rbcS and cbbX showed minimal rearrangement over minute timescales (Fig. S8). This observation aligns with recent data from the pennate diatom *P. tricornutum*, where *in vitro* phase-separated Rubisco demonstrates minimal rearrangement over similar timescales and the linker protein PYCO1 appears immobile *in vivo.*^51^ This lack of dynamic mixing opens a considerable question about how cbbX can sufficiently access inhibited Rubisco to reactivate it. Collectively, this indicates that the diatom pyrenoid has different mesoscale properties to the *C. reinhardtii* pyrenoid, and that once Rubisco is assembled into the pyrenoid there is minimal rearrangement. This observation aligns with those of the cyanobacterial carboxysome, where Rubisco forms ordered arrays upon packing.^43,51^

### Shell1 can self-assemble *in vitro* to form tubes and sheets

To form an encapsulating protein layer around the Rubisco matrix, we would expect that Shell proteins can undergo higher order self-assembly. To test this, we purified recombinantly-expressed Shell1 (Fig. S9A) from *E. coli* and performed cryogenic electron microscopy (cryo-EM). The formation of tubes and sheets (Fig. 6C and S9B) was observed in the micrographs. This observation is similar to the self-assembly of bacterial microcompartment hexamers, the main building block of carboxysome shells, which have been shown to self-assemble into both sheets^44,52^ and tubes *in vitro*.^47^ The ability to self-assemble into stable higher order structures likely explains the limited mobility of Shell1 observed *in vivo* and supports the hypothesis that Shell proteins could provide a continuous, encapsulating layer at the pyrenoid surface to function as a CO_2_ barrier, control metabolite flux, and/or provide structural integrity to the pyrenoid.

### Shell1/2 and Shell4 are essential for CCM function and pyrenoid architecture

The *in vitro* self-assembly of Shell1 into sheets and tubes and the immobile nature of Shell1 and Shell4 *in vivo* suggest they may be required for pyrenoid structural integrity. To test this, we used our MoClo Golden Gate system to simultaneously tag rbcS with mEGFP and disrupt either *Shell1/2* (*Shell1* and *Shell2)* or *Shell4* expression by CRISPR. Due to *Shell1* and *Shell2* having 93% DNA sequence similarity (Fig. S5C), single guide RNAs were designed to simultaneously target both genes (Fig. S10A). Edited lines were grown under high CO_2_ conditions and selected by mEGFP fluorescence. Biallelic gene editing was confirmed by Sanger sequencing (Fig. S10A,B). Imaging of rbcS-mEGFP in the *shell1/2* mutant revealed that cells lacking both *Shell1* and *Shell2* failed to form a lenticular pyrenoid and instead typically possessed a single spherical pyrenoid per chloroplast, although the presence of multiple pyrenoids was also observed (Fig. 6D). This suggests that Shell1/2 are required for the lenticular shape of the pyrenoid, and in their absence, the pyrenoid assembles into a sphere. Similarly, the *shell4* mutant failed to form a lenticular pyrenoid, forming more oval shaped pyrenoids that were subtly different in shape from the more spherical pyrenoids in the *shell1/2* mutant (Fig. 6D and S11). This hints that Shell4 may have a distinct structural role to Shell1/2, in line with their different localizations at the pyrenoid periphery. The more spherical appearance of Rubisco in the absence of the shell is consistent with surface tension effects observed in liquid-liquid phase separated condensates, suggesting dynamic Rubisco condensation may also have a role in pyrenoid matrix assembly *in vivo* in *T. pseudonana,* as shown *in vitro* for *P. tricornutum*^53^ and as seen for carboxysome assembly.^47^

Consistent with their disrupted pyrenoid morphology, the *shell1/2* and *shell4* mutants had significantly reduced growth at atmospheric CO_2_, which was fully rescued by supplying elevated CO_2_ (Fig. 6E), demonstrating that both Shell1/2 and Shell4 are required for a functional CCM. Interestingly, a *shell3* mutant did not show abnormal pyrenoid morphology by TEM (Fig. S10C and Fig. S12) and had no growth defect at ambient CO_2_ (Fig. 6E). Relative to other Shell homologs, Shell3 has a low abundance (Fig. 3D), and thus could potentially play a minor role or be compensated by Shell1/2, which have 73% amino acid sequence similarity. In a parallel study, it was shown that *shell1/2* knock-out mutants completely lack a protein shell around the pyrenoid.^47^ To see if our *shell4* mutant has a similar architectural defect, we performed cryo-electron tomography (cryo-ET).^47^ Wildtype *T. pseudonana* cells contain lenticular shaped pyrenoids, where the Rubisco matrix is encapsulated in a protein shell and bisected end-to-end by one or two PPTs that contain densities in their lumen (Fig. 6F).^47^ In contrast, the *shell4* mutant has misshaped pyrenoids with a diverse range of morphologies (Figs. 6G-I; S13). These include: 1) spherical pyrenoids that lack a visible shell and have abnormal PPTs that do not bisect the matrix (similar to the *shell1/2* mutant^27,53^); 2) pyrenoids with a shell that is detached from the Rubisco matrix; 3) pyrenoids that are encapsulated in a shell but have an atypical distorted shape and abnormal PPTs. Whole-cell proteomics on the *shell4* mutant showed the almost complete absence of Shell4 but largely unaltered levels of Shell1/2, suggesting that the Shell we observe in the *shell4* mutant is composed of Shell1/2 proteins (Fig. S14 and Table S7). Shells in the *shell4* mutants show apparent decreased affinity for the Rubisco matrix (detached shells) as well as apparent increased interaction with PPT membranes (attachment to PPTs located both within the matrix and displaced in the stroma). Interestingly, these mutant shells were also often observed to self-interact, forming double or triple layered sheets (Fig. S13). We therefore hypothesize that Shell4 promotes interaction with Rubisco, whereas Shell1/2 have high affinity for PPT membranes and Shell1/2 proteins in adjacent sheets. Taken together, the complete absence of a shell in the *shell1/2* mutant seen by cryo-ET^28,54^ and the presence of a shell but severely misshaped pyrenoids in the *shell4* mutant suggest that Shell1/2 and Shell4 have distinct roles: Shell1/2 form the main structural component of the shell, and Shell4 is critical for pyrenoid organization, ensuring that the Rubisco matrix, PPT, and shell are correctly assembled together. This is further supported by both the whole-cell proteomics, where Shell1/2 are more abundant than Shell4, as well as the pyrenoid interaction network, where Shell4 directly and indirectly is connected to several pyrenoid matrix components but Shell1 has no such connections.

The diverse pyrenoid structural defects in the *shell4* mutant makes it difficult to assign the high CO_2_ requiring phenotype to a specific pyrenoid structural defect. The disrupted PPTs proposed to be involved in inorganic carbon delivery to Rubisco could reduce inorganic carbon fluxes to the Rubisco matrix. If the shell has a role in minimizing CO_2_ diffusion, its disruption could make the pyrenoid leaky, thereby reducing the effective CO_2_ concentration around Rubisco. The shell could also be critical for efficient delivery and partitioning of Calvin cycle intermediates between the pyrenoid and the surrounding stroma. Finally, pyrenoid shape could be important to maximize membrane contact area and to minimize diffusion distances.

Collectively, these data support that the diatom Shell proteins 1, 2 and 4 are critical for pyrenoid architecture and CCM function, and may act as a CO_2_ diffusion barrier, although further experimental proof is required for the latter.

### Perspective

The development of a high-throughput tagging and APMS pipeline in a model diatom has enabled us to generate a spatial interaction network of the diatom pyrenoid, providing novel molecular insight into how diatoms help drive the global carbon cycle. We have identified and confirmed via localization 13 new pyrenoid components, of which a large number have no conserved functional domains. Six of these new components constitute a protein shell that encapsulates the pyrenoid and is found across diverse species with red plastids derived from secondary endosymbiotic events. Knock-out of the most abundant shell components, Shell1/2 and Shell4, resulted in large pyrenoid structural changes and poor growth at atmospheric levels of CO_2_. Cryo-ET on the *shell4* mutant indicates that Shell4 is critical for correct organization of the PPT, shell and matrix, whilst our data along with data in a parallel study^25,26^ indicates that Shell1/2 is most likely the major structural component of the shell.

Four additional pyrenoid components, DPC1-4, have no clear function that can be predicted from their sequence. DPC3 showed a shell-like localization pattern, co-localizing with Shell4 and interacting with both Shell4 and rbcS, suggesting a potential role in mediating shell-Rubisco matrix interactions. The molecular mechanism of Rubisco condensation to form the pyrenoid matrix is currently unknown in *T. pseudonana*, with no homologs or functional analogs of EPYC1 or PYCO1 (Rubisco likers in *C. reinhardtii* and *P. tricornutum*) identified in our study. This suggests that pyrenoid matrix formation may be different in *T. pseudonana*. The matrix localization and interaction partners of DPC2 and DPC4 suggest that they may have a central role in pyrenoid matrix assembly/function. DPC2, DPC3 and DPC4 are prime targets for future characterization.

Close to 50% of global carbon fixation is performed by biomolecular condensates of Rubisco. This includes prokaryotic cyanobacterial carboxysomes and eukaryotic algal pyrenoids. Nearly all of our data of pyrenoid structure and function comes from the green plastid lineage alga, *C. reinhardtii*. However, pyrenoids both between plastid lineages and within plastid lineages are proposed to have convergently evolved.^40^ Insights from our data suggest that diatom pyrenoids have similarities to both green plastid pyrenoids and prokaryotic carboxysomes. Similarities between the *T. pseudonana* and *C. reinhardtii* pyrenoid include dense Rubisco packaging around specialized thylakoid membranes (PPTs) for CO_2_ delivery, CO_2_ delivery to the PPTs via bestrophin family channels,^54^ and CO_2_ release within the acidic lumen driven by constrained localization of a carbonic anhydrase (θCA2; refs^55^ and this study) within the PPTs. However, the encapsulation by a protein shell layer composed of homologs, some with different sub-shell localizations, is analogous to carboxysome shell proteins.^47^ Additionally, the static nature of Rubisco, cbbX, and shell proteins contrasts with the dynamic properties of the *C. reinhardtii* pyrenoid and aligns more with carboxysomes. Another major outstanding question is the connection of the PPT with the broader thylakoid network; in both our TEM and cryo-ET, we are yet to visualize an unambiguous connection as seen in nearly all pyrenoid containing algae^7,23,24^ including *C. reinhardtii*^27^ and *P. tricornutum.*^56^ If there is no connection, this would align the *T. pseudonana* pyrenoid even closer to carboxysomes, and require a new functional model for the *T. pseudonana* CCM. It is tempting to postulate that the extension and diversification of the Shell protein family in *T. pseudonana* vs. *P. tricornutum* (6 homologs vs. 2 homologs; Fig. 4B) may be critical for the complete encapsulation of the Rubisco matrix and PPT, with Shell4 a strong candidate that could enable this.

In Figure 7, we propose two models for the *T. pseudonana* CCM, a more classical pyrenoid-based CCM where a PPT-thylakoid connection is present (Fig. 7A) and a completely Shell-encapsulated pyrenoid that would align closer to a carboxysome system (Fig. 7B). In both models, SLC4 family proteins contribute to sodium dependent HCO_3_^-^ transport at the plasma membrane.^57^ The mechanism of Ci transport across the four chloroplast membranes is still unknown, although SLC4 transporters are also proposed to play a role here^28^ along with the carbonic anhydrase LCIP63^25,26^ and the V-type ATPase.^58^ In the classical model, Ci delivery to the pyrenoid is potentially analogous to *C. reinhardtii*, relying on channeling of HCO_3_^-^ into the thylakoid lumen via BST1 and BST2.^59^ HCO_3_^-^ could then diffuse into the PPT, where it is dehydrated to CO_2_ via θCA2^60,61^ and diffuses to Rubisco packaged within the pyrenoid. In a more carboxysome-like system, HCO_3_^-^ accumulated in the stroma would have to diffuse across the shell and subsequently be dehydrated to CO_2_. CO_2_ release could potentially still occur via channeling into the PPT by bestrophins, and its release accelerated via θCA2. In the classical pyrenoid model, like in *C. reinhardtii*, the low luminal pH of the PPTs established by the light reactions of photosynthesis would provide protons for HCO_3_^-^ dehydration and the energetic driving force of the CCM.^62^ In a more carboxysome-like system, protons for HCO_3_^-^ dehydration could come from the Rubisco carboxylase reaction that has been modelled to produce two protons for every carboxylation reaction.^28^ In both models, CO_2_ leakage out of the pyrenoid would be minimized by the proteinaceous shell that encapsulates the pyrenoid. The shell is also critical for maintaining the correct architecture of the pyrenoid, including the lenticular shape of the pyrenoid that would minimize diffusion distances of CO_2_ from the PPTs to Rubisco.

**Figure 7:**
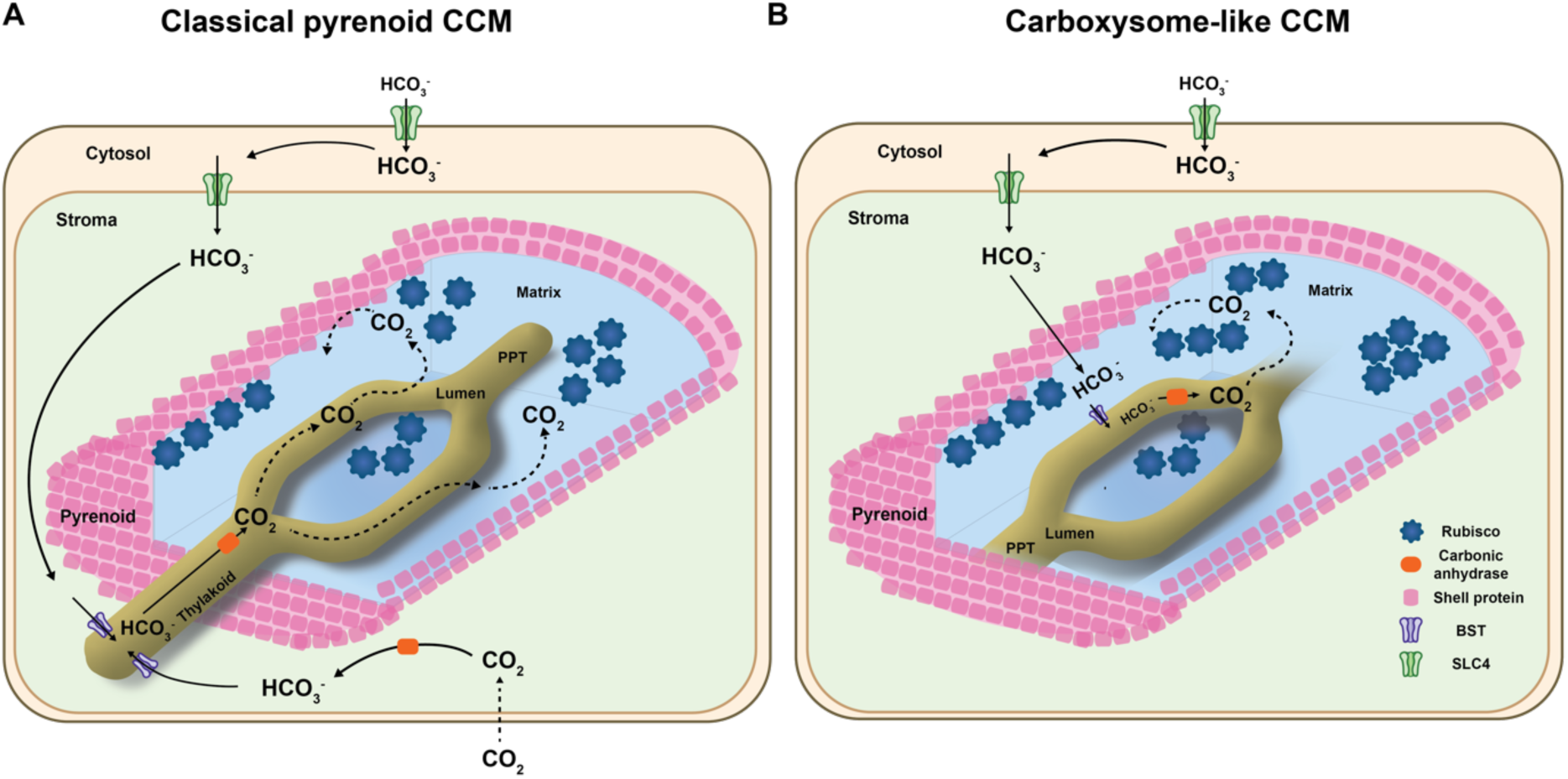
Structural and functional models of the *T. pseudonana* pyrenoid-based CO2-concentrating mechanism. B. Proposed model based on a classical pyrenoid CCM where there is a connection between the PPT and the broader thylakoid network. C. Proposed model based on a carboxysome-like CCM, where the shell completely encapsulates the pyrenoid and the PPT is separated from the broader thylakoid network.

Models integrate data from this study with the available literature. For simplicity, the multiple membranes encapsulating the chloroplast, and components with unknown or unclear function have been omitted. See text for further discussion.

Other than Rubisco, there appears to be no sequence or structural similarities between carboxysomes and *T. pseudonana* pyrenoid proteins. This further supports the convergent evolution of pyrenoids and that a broad range of biophysical, structural and functional properties, some previously associated with carboxysomes, can be expected as more pyrenoids are characterized across diverse alga.

A core structural component of pyrenoids is a CO_2_ leakage barrier, with the starch sheath in *C. reinhardtii* shown both experimentally and theoretically to be required for efficient CCM function.^17,18^ As engineering of a pyrenoid into plants progresses, a major future challenge will be CO_2_ diffusion barrier engineering.^17,22^ This is thought to require multiple starch synthesis-related steps correctly localized to the pyrenoid periphery. The diatom Shell proteins could potentially provide an alternative biotechnology solution to this challenge.

### Limitations of the Study

Well documented technical limitations of protein tagging could result in protein mislocalization and inaccurate protein-protein interactions. C-terminal tagging could result in masking interactions that could alter protein targeting and modulate native interactions with other proteins. The use of a constitutive promoter and expression of proteins in trans to native copies could also modify protein localization and interaction partners. Reported protein-protein interactions were not validated by a parallel method and may not mean a direct interaction but could be mediated through an additional component. Although DPC3 and Shell4 were shown to interact and colocalize, colocalization does not confirm a direct interaction.

Where possible we validated two independently tagged lines for localization, however in some cases only a single stable line was generated. For *shell4* mutants, four independent lines were sequence verified for biallelic editing. Both pyrenoid shape disruption by rbcs-mEGFP imaging and growth defects at ambient CO_2_ were seen across all edited lines, and a single representative line was chosen for further detailed studies. For *shell1/2* mutants, multiple lines were generated that resulted in pyrenoid shape disruption by rbcs-mEGFP imaging, however only one was confirmed for biallelic editing of both *Shell1* and *Shell2*; this line was prioritized for further studies.

Whilst we propose that the shell may act as a CO_2_ barrier analogous to the carboxysome shell, we provide no direct evidence for this. We also do not understand how the Rubisco substrate, ribulose 1,5-bisphosphate (RuBP), and product, 3-phosphoglycerate (PGA), cross the shell. Although we show that Shell1 and Shell4 do not colocalize, we do not understand how the other Shell components are orientated relatively to each other, whether Shell proteins other than Shell1 can homo-oligomerize, and whether Shell proteins can hetero-oligomerize. All our images are snapshots of living cells. Proteins could change localization and interactions depending on cell-cycle state and during pyrenoid division. Further, the dynamics of Shell1, Shell4, Rubisco, and cbbX may change at different states of the cell cycle to enable pyrenoid division and pyrenoid growth. Finally, how the pyrenoid matrix is assembled is still an open question.

## Materials and methods

### Strains and Culturing

The background *Thalassiosira pseudonana* strain for all experiments was wildtype (WT) CCAP1085/12 (Scottish Culture Collection of Algae and Protozoa, equivalent to CCMP1335). WT cells were axenically maintained in artificial sea water (ASW) (32 g L^-1^, Instant Ocean SS15-10) supplemented with half-strength (F/2) Guillard F solution^64^ at 20°C under continuous illumination of ∼50 μmol photons m^-2^ s^-1^. All strains were grown at ambient CO_2_ except *shell1/2* deletion, *shell4* deletion and Shell1-mEGFP lines, which were maintained under 1% CO_2_. Exponentially growing cultures were used for growth assays. Once the pre-cultures reached 2-4 x 10^6^ cell mL^-1^, cells were harvested by centrifugation at 3000 x*g* for 10 min. Then the cells were resuspended at a concentration of 3 x 10^5^ cell mL^-1^ in 10 mL of fresh media. To ensure optimal gas exchange, cultures were grown in 6-well plates swirling twice a day and incubated in growth chambers with water-saturated 0.04% CO_2_ (ambient air, LC) or 1% CO_2_ (HC) in air. Cell density was monitored daily by counting cells using a digital haemocytometer (Countess II FL, Life Technology). All experiments were performed in biological triplicates.

### Episome Assemblies using Golden Gate Cloning

Level 0, 1, and 2 (L0, L1, and L2) plasmids were assembled by Golden Gate (GG) cloning^65^ using the custom parts from the diatom MoClo framework.^35^ Using genome version ASM14940v2, target genes without stop codons were synthesized by Twist Bioscience (Table S4). The regulatory elements, fucoxanthin chlorophyll a/c-binding protein (FCP) promoter and terminator, and the fluorescent protein (FP) tags (mEGFP and mScarlet-I) harboring L0 plasmids were used to build L1 plasmids together with the gene-of-interest (GOI) for FP tagging. Subsequently, L1 plasmids were assembled into L2 plasmids (episomes) in the following order: episomal maintenance elements (position 1, P1),^65^ nourseothricin (NAT) resistance cassette (P2), FP tagged GOI (P3), and for dual-FP tagging (P4). For the CRISPR-Cas9 knock-out episomes, Cas9 occupied P3 and two sgRNAs for each target both under the U6 promoter occupied P4 and P5. P6 was used for simultaneous knock-out and rbcS-mEGFP tagging. Each GG assembly was performed in 20 μL containing 40 fmols of each component with 10x ligase buffer (NEB), 10 units T4 DNA ligase (NEB), and 10 units restriction enzyme (BsaI for L1 assembly or BpiI for L0/L2 assembly, Thermo Fisher Scientific). The reaction was incubated in a thermocycler by switching between 37°C and 16°C for 5 min intervals in a total of 20-30 cycles, followed by 37°C for 5 min and terminated after incubating at 65°C for 20 min. 3 μL of the reaction were transformed into 50 μL chemically competent DH5α *E. coli* cells.

### *T. pseudonana* Transformation via Bacterial Conjugation

Episomes were delivered to *T. pseudonana* via bacterial conjugation according to^66^ with minor modifications. Episome plasmids were transformed into *E. coli* (TransforMax EPI300) harboring the pTA_Mob^67^ mobility plasmid (gift from R. Lale) via electroporation (Bio-Rad). Transformed cells were spread onto LB agar plates containing both gentamycin (10 μg mL^-1^) and kanamycin (25 μg mL^-1^) for selection overnight at 37°C. Colonies were inoculated for subsequent conjugation. Cultures (150 mL) grown at 37°C to OD600 of 0.3-0.4 were harvested by centrifugation (3,000 x*g*, 5 min) and resuspended in 800 μL of SOC media. Liquid grown *T. pseudonana* WT culture was harvested by centrifugation (3,000 x*g*, 5 min) and resuspended at a concentration of 2x10^8^ cells mL^-1^ in ½ASW-F/2. Equal volume (200 μL) of *E. coli* and *T. pseudonana* WT cells were gently mixed by pipetting. Next the mixture of cells was plated on ½ASW-F/2, 5% LB, 1% agar plates and incubated in the dark for 90 min at 30°C. The plates were transferred to 20°C with continuous illumination (∼50 μmol photons m^-2^ s^-1^) and grown overnight. Next day, 500 μL of ½ASW-F/2 medium was added to the plate for scraping and resuspending the cells. Up to 200 μL of resuspended cells were spread onto 1% (w/v) ½ASW-F/2 agar plates with 100 μg mL^-1^ nourseothricin for selection. Colonies appeared after 6-14 days.

### Fluorescence Screening by Flow Cytometry

mEGFP and mScarlet-I expression was analyzed by flow cytometry using either CytoFLEX LX355 or 375 (Beckman Coulter) analyzers. Forward scattered (FSC) and side scattered photons by the 488 nm laser were used to distinguish diatoms from cell culture debris. FSC-height versus FSC-area signal was used to separate single events from sample aggregates. Chlorophyll autofluorescence excited by 561 nm laser and emitted photons detected with 675/25 filter was used to ensure all the diatom cells were fully intact. mEGFP fluorescence excited by the 488 nm laser was detected by an avalanche photodiode detector with 525/40 bandpass filter. All the data analysis was done using CytExpert software (Beckman Coulter).

### Microscopy

Fluorescence imaging was performed using a Zeiss LSM880 confocal microscope with a 63x objective, 1.4 numerical aperture (NA) Plan-Apo oil-immersion lens (Carl Zeiss). All imaging was done in Airyscan mode except for the rbcS-mEGFP line, which was imaged in confocal mode. 20 μL of cell suspension were pipetted on 8 well μ-Slide chambered coverslips (ibidi) overlaid with 180 μL of 1.5% F/2-low-melting point agarose (Invitrogen) for imaging. Excitation lasers and emission filters were as follows: mEGFP excitation 488 nm, emission 481-541 nm; mScarlet-I excitation 561 nm, emission 561-633 nm; and chlorophyll excitation 633 nm, emission 642-712 nm. All the microscopic images were processed using Fiji.^68^

### Co-immunoprecipitation and affinity purification

For rbcL coIP, 50 mL of WT *T. pseudonana* cells grown in log phase (2-3 x 10^6^ cells mL^-1^) were harvested by centrifugation (3,000 x*g*, 10 min). The pellets were resuspended in CoIP buffer (20 mM Tris-HCl pH 8.0, 50 mM NaCl, 0.1 mM EDTA, 12.5% glycerol) containing 5 mM DTT and protease inhibitor cocktail tablets (PIs, cOmplete EDTA-free, Roche). Cells were lysed by sonication for 3 min (ON 3 sec, OFF 12 sec). The lysates were centrifuged for 20 min (20,817 x*g*, 4°C) to separate the supernatant (soluble lysate) from the pellet. 200 μL of Protein A (Dynabeads Protein A, Invitrogen) beads were washed twice in coIP buffer containing PIs. 32 μg of anti-rbcL antibody in 500 μL coIP buffer containing PIs was added to the washed beads and incubated at 4°C for 2 hours. After incubation beads were washed twice in coIP buffer containing PIs. For blocking, 500 μL of BSA (2 mg mL^-1^) was added and incubated at 4°C for 1 hour. After incubation beads were washed twice in coIP buffer containing PIs. Subsequently, the soluble lysates were added to protein A beads primed with antibody and incubated at 4°C for 3 hours. After incubation, beads were washed three times with coIP buffer containing PIs and 0.1% digitonin (SigmaAldrich). For elution, 200 μL of 1x SDS loading dye was added and boiled at 95°C for 5 min. The supernatant was collected without any beads and ran on an SDS-PAGE gel for ∼1.5 cm. Gels were sliced for further in-gel digestion for LC-MS/MS (see below).

For mEGFP tagged lines AP, 50 mL of GFP tagged *T. pseudonana* lines grown in exponential phase (2-3 x 10^6^ cells mL^-1^) were harvested by centrifugation (3,000 x*g*, 10 min). The pellets were resuspended in an immunoprecipitation (IP) buffer (200 mM D-sorbitol, 50 mM HEPES, 50 mM KOAc, 2 mM Mg(OAc)_2_, 1 mM CaCl_2_) containing protease inhibitor cocktail tablets (cOmplete EDTA-free, Roche), 2% digitonin (SigmaAldrich), 1mM PMSF, 0.5 mM NaF and 0.15 mM Na_3_VO_4_. Cells were lysed by sonication for 30 sec (On 3 sec, Off 15 sec) twice. The lysates were centrifuged for 20 min (20,817 x*g*, 4°C) and the supernatant was incubated with mEGFP-Trap Agarose beads (ChromoTek) for 1 hour according to the manufacturer’s instructions. Subsequently, beads were washed twice with an IP buffer containing 0.1% digitonin and a final wash without digitonin. All steps were performed at 4°C.

### Mass Spectrometry

For rbcL coIPMS, samples were in-gel digested with 0.2 μg Sequencing-grade, modified porcine trypsin (Promega), following reduction with 1.5 mg ml^-1^ dithioerythritol and alkylation with 9.5 mg mL^-1^ iodoacetamide. Digests were incubated overnight at 37°C. Peptides were extracted by washing three times with aqueous 50% (v:v) acetonitrile containing 0.1% (v:v) trifluoroacetic acid, before drying in a vacuum concentrator and reconstituting in aqueous 0.1% (v:v) trifluoroacetic acid. Peptides were loaded onto an mClass nanoflow UPLC system (Waters) equipped with a nanoEaze M/Z Symmetry 100 Å C_18_, 5 µm trap column (180 µm x 20 mm, Waters) and a PepMap, 2 µm, 100 Å, C 18 EasyNano nanocapillary column (75 mm x 500 mm, Thermo). The trap wash solvent was aqueous 0.05% (v:v) trifluoroacetic acid and the trapping flow rate was 15 µL min^-1^. The trap was washed for 5 min before switching flow to the capillary column. Separation used gradient elution of two solvents: solvent A, aqueous 0.1% (v:v) formic acid; solvent B, acetonitrile containing 0.1% (v:v) formic acid. The flow rate for the capillary column was 330 nL min^-1^ and the column temperature was 40°C. The linear multi-step gradient profile was: 3-10% B over 5 mins, 10-35% B over 85 mins, 35-99% B over 10 mins and then proceeded to wash with 99% solvent B for 5 min. The column was returned to initial conditions and re-equilibrated for 15 min before subsequent injections. The nanoLC system was interfaced with an Orbitrap Fusion Tribrid mass spectrometer (Thermo) with an EasyNano ionisation source (Thermo). Positive ESI-MS and MS2 spectra were acquired using Xcalibur software (version 4.0, Thermo). Instrument source settings were: ion spray voltage, 1900-2100 V; sweep gas, 0 Arb; ion transfer tube temperature; 275°C. MS 1 spectra were acquired in the Orbitrap with: 120,000 resolution, scan range: *m/z* 375-1,500; AGC target, 4e^5^; max fill time, 100 ms. Data dependent acquisition was performed in top speed mode using a 1 s cycle, selecting the most intense precursors with charge states >1. Easy-IC was used for internal calibration. Dynamic exclusion was performed for 50 s post precursor selection and a minimum threshold for fragmentation was set at 5e^3^. MS2 spectra were acquired in the linear ion trap with: scan rate, turbo; quadrupole isolation, 1.6 *m/z*; activation type, HCD; activation energy: 32%; AGC target, 5e^3^; first mass, 110 *m/z*; max fill time, 100 ms. Acquisitions were arranged by Xcalibur to inject ions for all available parallelizable time. Tandem mass spectra peak lists were extracted from Thermo .raw files to .mgf format using MSConvert (ProteoWizard 3.0). Mascot Daemon (version 2.6.0, Matrix Science) was used to submit searches to a locally-running copy of the Mascot program (Matrix Science Ltd., version 2.7.0). Peak lists were searched against the *Thalassiosira pseudonana* subsets of UniProt and NCBI with common proteomic contaminants appended. Search criteria specified: Enzyme, trypsin; Max missed cleavages, 2; Fixed modifications, Carbamidomethyl (C); Variable modifications, Oxidation (M); Peptide tolerance, 3 ppm; MS/MS tolerance, 0.5 Da; Instrument, ESI-TRAP. Peptide identifications were collated and filtered using Scaffold (5.2.0, Proteome Software Inc). Peptide identifications were accepted if they could be established at greater than 51.0% probability to achieve an FDR less than 1.0% by the Percolator posterior error probability calculation. Protein identifications were accepted if they could be established at greater than 5.0% probability to achieve an FDR less than 1.0% and contained at least 2 identified peptides.

For mEGFP tagged lines APMS, samples were on-bead digested using Chromotek’s recommended procedure for NanoTraps: protein was digested overnight at 37°C with 25 μL 50 mM Tris-HCl pH 7.5, 2 M urea, 1mM DTT, 5 μg ml^-1^ Sequencing Grade Modified Trypsin (Promega). Peptides were eluted with 50 mM Tris-HCl pH 7.5, 2 M urea, 5 mM iodoacetamide before loading onto EvoTip Pure tips for desalting and as a disposable trap column for nanoUPLC using an EvoSep One system. A pre-set EvoSep 60 SPD gradient was used with a 8 cm EvoSep C_18_ Performance column (8 cm x 150 μm x 1.5 μm). The nanoUPLC system was interfaced to a timsTOF HT mass spectrometer (Bruker) with a CaptiveSpray ionisation source (Source). Positive PASEF-DDA, ESI-MS and MS2 spectra were acquired using Compass HyStar software (version 6.2, Thermo). Instrument source settings were: capillary voltage, 1,500 V; dry gas, 3 l min-1; dry temperature; 180°C. Spectra were acquired between *m/z* 100-1,700. The following TIMS settings were applied as: 1/K0 0.6-1.60 V.s cm-2; Ramp time, 100 ms; Ramp rate 9.42 Hz. Data dependent acquisition was performed with 10 PASEF ramps and a total cycle time of 1.17 s. An intensity threshold of 2,500 and a target intensity of 20,000 were set with active exclusion applied for 0.4 min post precursor selection. Collision energy was interpolated between 20 eV at 0.5 V.s cm^-2^ to 59 eV at 1.6 V.s cm^-2^. Pick picking, database searching, significance thresholding and peak area integration was performed using FragPipe (version 19.1). Data were searched against UniProt reference proteome UP000001449, appended with common contaminants and concatenated with reversed sequences for false discovery calculation. Search criteria specified: Enzyme, trypsin; Max missed cleavages, 2; Fixed modifications, Carbamidomethyl (C); Variable modifications, Oxidation (M), Acetylation (Protein N-term); Peptide tolerance, 10 ppm; MS/MS tolerance, 10 ppm; Instrument, IM-MS. Peptide identifications were filtered using Percolator and ProteinProphet to 1% PSM FDR, protein probabilities >99%, best peptide probability >99% and a minimum of two unique peptides. Peak area quantification was extracted using IonQuant with match between run applied. Feature detection tolerances were set to: MS1 mass <10 ppm; RT < 0.4 min; and IM (1/k0) <0.05.

For whole-cell MS samples were ran in triplicate. 50 mL of WT cells were grown in ambient CO_2_ conditions and harvested at the log phase (2-3 x 10^6^ cells mL^-1^) by centrifugation (3,000 x*g*, 10 min). Shell4 mutant lines were grown in 1% CO_2_. Cells were lysed by sonication (3 min: ON 3 s, OFF 12 s) in CoIP buffer containing 5 mM DTT and PIs followed by centrifugation for 20 min (20,817 x*g*, 4°C). Supernatant was run on an SDS-PAGE gel for ∼1.5 cm. In-gel digestion was performed with the addition of 0.2 mg sequencing-grade, modified porcine trypsin (Promega), following reduction with dithioerythritol and alkylation with iodoacetamide. Digests were incubated overnight at 37°C. Peptides were extracted by washing three times with aqueous 50% (v:v) acetonitrile containing 0.1% (v:v) trifluoroacetic acid, before drying in a vacuum concentrator and reconstituting in aqueous 0.1% (v:v) trifluoroacetic acid.

S-Trap™ micro spin column (PROFITI) digestions were performed using the manufacture’s protocol. Briefly, protein in 5% SDS (w:w) lysis buffer was reduced with tris(2-carboxyethyl)phosphine and alkylated with methyl methanethiosulfonate before acidification with phosphoric acid and dilution into 100 mM triethylammonium bicarbonate (TEAB) in 90% (v:v) methanol. Protein was washed on s-Trap with the same buffer five times before digestion for 2 h at 47°C with 2 mg Promega Trypsin/Lys-C mixed protease in 50 mM TEAB.

Peptides were loaded onto EvoTip Pure tips for nanoUPLC using an EvoSep One system. A pre-set 30SPD gradient was used with a 15 cm EvoSep C_18_ Performance column (15 cm x 150 mm x 1.5 mm).

The nanoUPLC system was interfaced to a timsTOF HT mass spectrometer (Bruker) with a CaptiveSpray ionisation source. Positive PASEF-DIA, nanoESI-MS and MS^2^ spectra were acquired using Compass HyStar software (version 6.2, Bruker). Instrument source settings were: capillary voltage, 1,500 V; dry gas, 3 l/min; dry temperature; 180°C. Spectra were acquired between m/z 100-1,700. DIA windows were set to 25 Th width between m/z 400-1201 and a TIMS range of 1/K0 0.6-1.60 V.s/cm2. Collision energy was interpolated between 20 eV at 0.65 V.s/cm2 to 59 eV at 1.6 V.s/cm2.

LC-MS data, in Bruker .d format, was processed using DIA-NN (1.8.2.27) software and searched against an *in silico* predicted spectral library, derived from the *Thalassiosira pseudonana* subset of UniProt (11934 protein sequences). Search criteria were set to maintain a false discovery rate (FDR) of 1% with heuristic protein inference. High-precision quant-UMS was used for extraction of quantitative values within DIA-NN. Peptide-centric output in .tsv format, was pivoted to protein-centric summaries using KNIME 5.1.2 and data filtered to require protein q-values < 0.01 and a minimum of two peptides per accepted protein. Raw proteomic mass spectrometry data and search results files are referenced in ProteomeXchange (PXD052522) and can be accessed via MassIVE (MSV000094846) (doi:10.25345/C53J39C2Q).

### Protein-protein interaction network analysis

Protein abundances quantified using MS2 spectral count measurements of fragment ions from all sample triplicates were run through a CompPASS package in R Studio (https://github.com/dnusinow/cRomppass/blob/master/R/cRomppass.R) and a control IP inclusive variation of SAINT analysis in Ubuntu using standard parameters.^69^ The WD and AvgP scores respectively generated were used as measures of interaction strength between bait and prey proteins. Only interactions which fell in both the top 2.2% WD score and 1% AvgP score were filtered as high confidence interactors. Prior to analysis bait spectral count data was set to zero to minimize data skewing due to the typically high spectral counts and the inability to distinguish between mEGFP tagged bait and untagged native protein.

### *In vivo* fluorescence recovery after photobleaching

FRAP experiments were performed using a Zeiss LSM980 confocal microscope with a 63x objective 1.4 numerical aperture (NA) Plan-Apo oil-immersion lens (Carl Zeiss). Samples were prepared as in confocal microscopy and overlaid with 200 μL of ibidi anti-evaporation oil. 20 pre-bleach images were taken prior to bleaching (60% 488 nm intensity, 1 cycle). All the images were processed by Fiji.^70^ The Image Stabilizer plugin (4 pyramid levels, 0.99 template update coefficient) output of the brightfield images were used to stabilize the fluorescence images. The mean gray values were measured for the bleached, unbleached and background ROIs. Background values were subtracted from bleached and unbleached values before photobleach normalization using the unbleached reference was completed. The average pre-bleach intensity was used for full-scale normalization.

### Data Visualization

Network visualization of the interaction network was done using Cytoscape (version 3.10.0) (https://cytoscape.org/). The phylogenetic tree was done in Geneious Prime (2023.2.1). Adobe Illustrator (2023), Excel (Microsoft) and Prism 10 (GraphPad) were used to generate the figures.

### AlphaFold Structure Prediction

AlphaFold structure predictions for Shell 1-6 and DPC3 without signal peptide and chloroplast transit peptides were done using ColabFold v1.5.2-patch: AlphaFold2 using MMseqs2 (https://colab.research.google.com/github/sokrypton/ColabFold/blob/main/AlphaFold2.ipynb).

### Protein model for Rubisco

*Thalassiosira antarctica* Rubisco model (PDB: 5MZ2) was used to predict the peptide regions utilized for raising the anti-rbcL antibody using ChimeraX.^71^

### Transmission Electron Microscopy

50 mL of cell culture grown in log phase were harvested by centrifugation (3,000 x*g*, 5 min). Cells were fixed in 0.1 M cacodylate buffer (SigmaAldrich), pH 7.4, containing 2.5% glutaraldehyde (SigmaAldrich), 2% formaldehyde (Polysciences) for 1 hour at room temperature. They were then rinsed twice in phosphate-buffered saline (PBS, pH 7.4) for 15 min by removing the liquid at the top of the tube and leaving the cells undisturbed at the bottom of the tube. The cells were secondary fixed with Osmium Tetroxide (1%, in buffer pH 7.2, 0.1M) for 1 hour. After rinsing twice with buffer, again removing the liquid above the cells whilst leaving the cells undisturbed, the cells were dehydrated through an ethanol series of 30, 50, 70, 90 and 100%, with each rinse for 15 min. To ensure thorough dehydration the 100% ethanol step was repeated 3 times. The ethanol was then replaced by Agar low viscosity resin by placing it in increasing concentrations of resin (30% resin: 70 % ethanol. 50:50, 70:30 each change was left for at least 12 hours) until it was in 100% resin. To ensure complete resin infiltration, the 100% resin step was repeated 3 times leaving it overnight between changes. The Eppendorf tubes were then placed in an embedding oven and the resin polymerized at 60°C overnight. The resulting blocks were sectioned with a Leica Ultracut E ultra microtome using a diatome diamond knife. The sections were then stained using a saturated solution of Uranyl acetate (for 15 min) and Reynold’s Lead citrate (15 min). The sections were imaged using a JEOL 1400 TEM.

### Blast search

Full amino acid sequence of Shell 1-6 was used for Blastp with default settings: ’Standard databases’, ‘Non-redundant protein sequences (nr)’, ‘blastp (protein-protein BLAST)’ ‘Max target sequences=100’ ‘Word size =5’ ‘ Max matches in a query range = 0’ ‘Matrix = BLOSUM62’ ‘Gap Costs= Existence: 11 Extension: 1’ ‘Compositional adjustments = Conditional compositional score matrix adjustment’ and ‘Expect threshold =1’. Hits were sorted by highest to lowest alignment length, and a cut-off length of 100 amino acids was employed.

### Phylogeny

The phylogenetic tree was build using Geneious Prime (2023.2.1). 136 proteins were used for MAFFT alignment using default settings (Algorithm = ‘Auto’, Scoring matrix = ‘BLOSUM62’, Gap open penalty = ‘1.53’, Offset value = ‘0.123’). Phylogenetic tree analysis was performed using RAxML with the following settings: Protein Model = ‘GAMMA BLOSUM62’, Algorithm = ‘Rapid Bootstrapping’, Number of starting trees or bootstrap replicates = ‘1000’, Parsimony random seed = ‘1’. A consensus tree was generated by selecting Consensus Tree Builder, Create Consensus Tree, Support Threshold % = 0, Topology Threshold % = 0, Burn-in % = 0, Save tree(s) separately. *C. reinhardtii* BST1-3 sequences (Cre16.g662600, BST1; Cre16.g663400, BST2; Cre16.g663450, BST3) were added to root the tree.

### Shell1 purification

A modified pOPT expression plasmid^72^ containing *shell1* with N-terminal His6-MBP tag and C-terminal GST tag was transformed into *E. coli* expression strain C41 DE3. Cells were grown in 2 L of liquid LB medium with 100 μg mL^-1^ carbenicillin until OD_600_ reached 0.6 when the culture was induced with 1 mM IPTG for 6 h. Harvested cells were resuspended in a lysis buffer. The lysis buffer contained 50 mM Tris-HCl, pH 8.0, 1x cOmplete EDTA-free protease inhibitor with 500 mM NaCl and 2 mM PMSF. The cells were lysed by cell disruption (30 kPSI) and the lysate cleared by 10 min of centrifugation at 7,000 x*g* and 30 min at 20,000 x*g* before being filtered with a 0.2 μm filter. The supernatant was applied onto a 5 ml HisTrap Fast Flow nickel affinity column (Cytiva) equilibrated with buffer A (50 mM Tris-HCl pH 8.0, 500 mM NaCl) followed by 4CV wash with lysis buffer. The bound protein was then eluted using a linear gradient of imidazole up to 500 mM concentration. The elution was then treated with TEV protease and 3C protease while dialysed overnight at 4°C into 50 mM Tris-HCl, pH 8.0, 500 mM NaCl buffer. The sample was then applied to both a HisTrap Fast Flow nickel column and a GSTrap column to remove the tags. The Shell protein remained in the flowthrough. The protein was then concentrated using an Amicon Ultra concentrator 10,000 MW, applied onto a Superdex 200 pg 16/600 Column equilibrated with buffer A to further ensure the purity of the sample, and concentrated again with an Amicon Ultra Concentrator if needed.

QUANTIFOIL^®^ R 1.2/1.3 grids were glow discharged with PELCO easiGlow system using a pressure of 0.26 mBar, for 60 seconds at a 20 mA current. Shell1 protein sample in 50 mM Tris-HCl (pH 8.0), 50 mM NaCl at 2.3 mg mL^-1^ concentration was applied onto the grid, blotted for 8 s using force 10 with a FEI Mark IV Vitrobot (ThermoFisher Scientific) in a chamber at 4°C and 95% relative humidity. Grids were then rapidly plunge frozen in liquid ethane.

Data was collected on a 200 kV Glacios cryo-electron microscope with a Falcon IV direct electron detector at the University of York. Automated data collection was performed using EPU in AFIS mode (ThermoFisher Scientific). Micrographs were collected at a nominal magnification of 120,000x and total electron dose of 50 e^-^/Å^2^ with pixel size of 1.2 Å^2^. Total exposure was 8 s. The 100 μm objective aperture was inserted and the C2 aperture was 50 μm. Defocus values were: -2.0, -1.6, -1.4, -1.2, -1.0.

### Cellular cryo-electron tomography

*T. pseudonana* cultures were grown as described above, with ΔtpPyShell4 being supplemented with 5 mM NaCO3 in the F/2 medium. Cells were sedimented at 800 x*g* for 5 min prior to vitrification, 4 μL of cell suspension was applied to 200-mesh R2/1 carbon-film covered copper grids (Quantifoil Micro Tools), and grids were blotted and plunge-frozen using a Vitrobot Mark IV (Thermo Fisher Scientific). EM grids were clipped into Autogrid supports (Thermo Fischer Scientific) and loaded into Aquilos 2 FIB-SEM instruments (Thermo Fischer Scientific), where they were thinned with a Gallium ion beam as previously described.^73^ The resulting EM grids with thin lamellae were transferred to a transmission electron microscope for tomographic imaging. Datasets were acquired on three different microscopes: a 300 kV Titan Krios G4i microscope (Thermo Fisher Scientific), equipped with a Selectris X post-column energy filter (Thermo Fisher Scientific) and a Falcon 4 direct electron detector (Thermo Fisher Scientific) (“M1”); a 300 kV Titan Krios G3i microscope (Thermo Fisher Scientific) equipped with a BioQuantum post-column energy filter (Gatan) and a K3 direct electron detector (Gatan) (“M2”); and Titan G4 equipped with a monochromator, a Selectris X energy filter, and a Falcon 4 camera (“M3”). Tilt-series were obtained using SerialEM 3.8 software^74^ or Tomo5 (Thermo Fisher Scientific). In all cases, tilt-series were acquired using a dose-symmetric tilt scheme,^75^ with 2° steps totaling 60 tilts per series. Each image was recorded in counting mode with ten frames per second. For M1 and M3, data was acquired in EER mode. The target defocus of individual tilt-series ranged from −2 to −5 µm. Total dose per tilt series was approximately 120 e-/Å2. Image pixel sizes for microscopes M1, M2, and M3 were 2.42 Å, 2.143/2.685 Å, and 2.38 Å, respectively.

Tomograms were subsequently reconstructed. First, TOMOMAN Matlab scripts (version 0.6.9 https://github.com/williamnwan/TOMOMAN/; https://doi.org/10.5281/zenodo.4110737)^76^ were used to preprocess the tomographic tilt series data. Raw frames were then aligned using MotionCor2 (version 1.5.0)^77^ before dose-weighting the tilt-series series,^66^ followed by manual removal of bad tilts. The resulting tilt-series (binned 4 times) were aligned in IMOD (version 4.11)^74^ using patch tracking and were reconstructed by weighted back projection. Cryo-CARE (version 0.2.1)^75^ was applied on reconstructed tomogram pairs from odd and even raw frames to enhance contrast and remove noise. Snapshots of denoised tomograms were captured using the IMOD 3dmod viewer. Denoised tomograms were used as input for automatic segmentation using MemBrain.^76^ The resulting segmentations were manually curated in Amira (version 2021.2; Thermo Fisher Scientific). Rubisco enzymes were template matched using PyTOM (version 0.0.6)^77^ on bin4 tomograms. The resulting segmentations and particle coordinates were visualized in ChimeraX.^66^

## Data availability

All mass spectrometry and proteomic identification data are referenced in ProteomeXchange (PXD052522) and can be accessed via MassIVE (MSV000094846).

## Author contributions

O.N. and L.C.M.M. designed and supervised the study. O.N. carried out the experiments unless otherwise mentioned. M.D. and B.E. performed cryo-ET of the *shell4* mutant. S.M. purified the Shell1 protein and carried out cryo-EM with support from J.N.B. A. M. and O.N. performed the phylogenetic analysis of Shell proteins. A.D., M.D. and J.B. provided bioinformatics and data analysis support. A.D. oversaw the mass spectrometry and peptide mapping. O.N., C.M., and L.C.M.M. analyzed and interpreted the data. O.N. created the figures. O.N. and L.C.M.M. wrote the manuscript with input from all authors.

## Acknowledgements

We thank Oliver Mueller-Cajar for kindly sharing the anti-rbcL antibody, the Technology Facility in the York Department of Biology for the access/support for microscopy and flow cytometry, and members of the Mackinder Lab for fruitful discussions. Thanks to Glenn Harper for optimizing the TEM. Cryo-ET was performed at the University of Basel using instrumentation provided by the BioEM core facility and computation provided by the sciCORE high-performance computing center. The York Centre of Excellence in Mass Spectrometry was created thanks to a major capital investment through Science City York, supported by Yorkshire Forward with funds from the Northern Way Initiative, and subsequent

---

**Figure S1.**
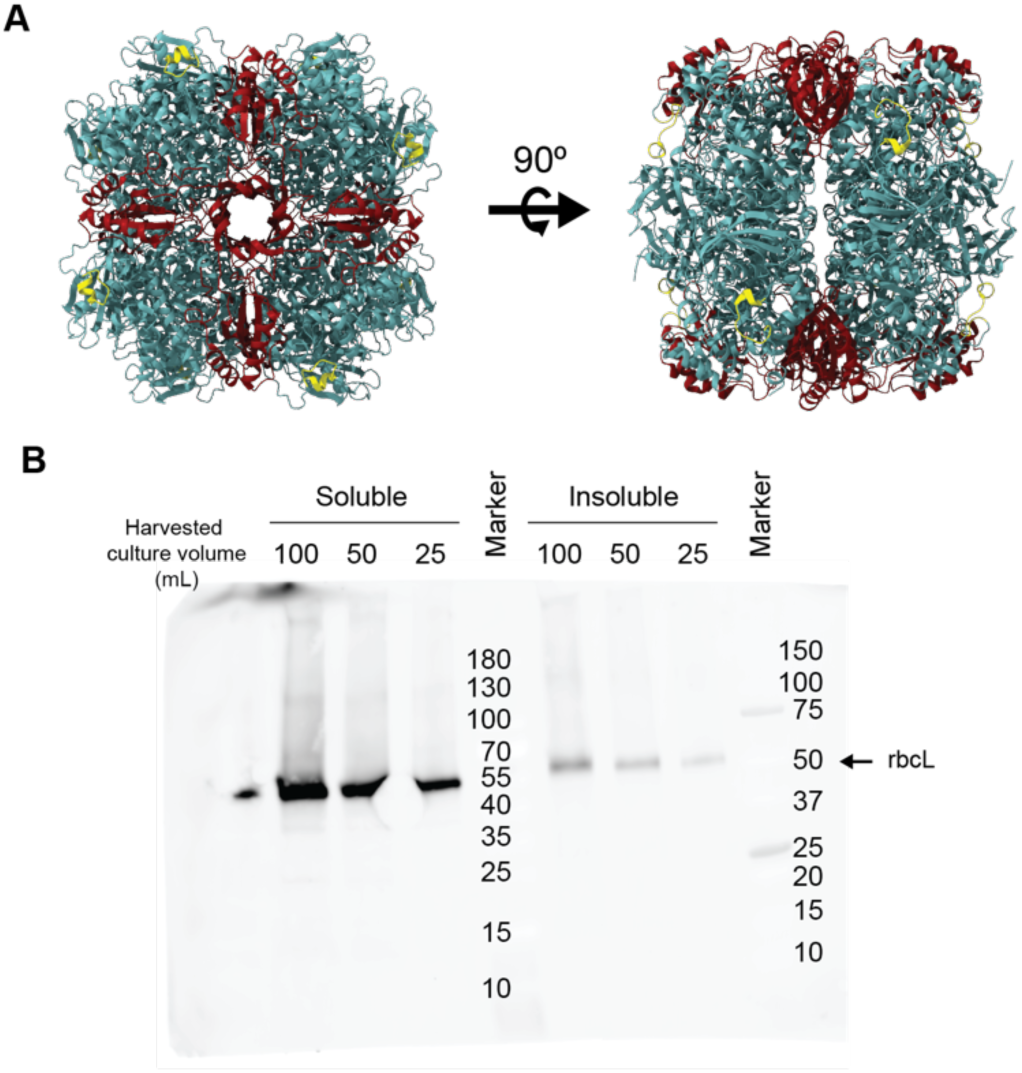
RbcL antibody. A. An antibody was raised against a surface exposed region (yellow) of the rbcL (green). RbcS is shown in red. B. Immunoblot using the rbcL antibody against the soluble and insoluble cell fractions of *T. pseudonana*.

**Figure S2.**
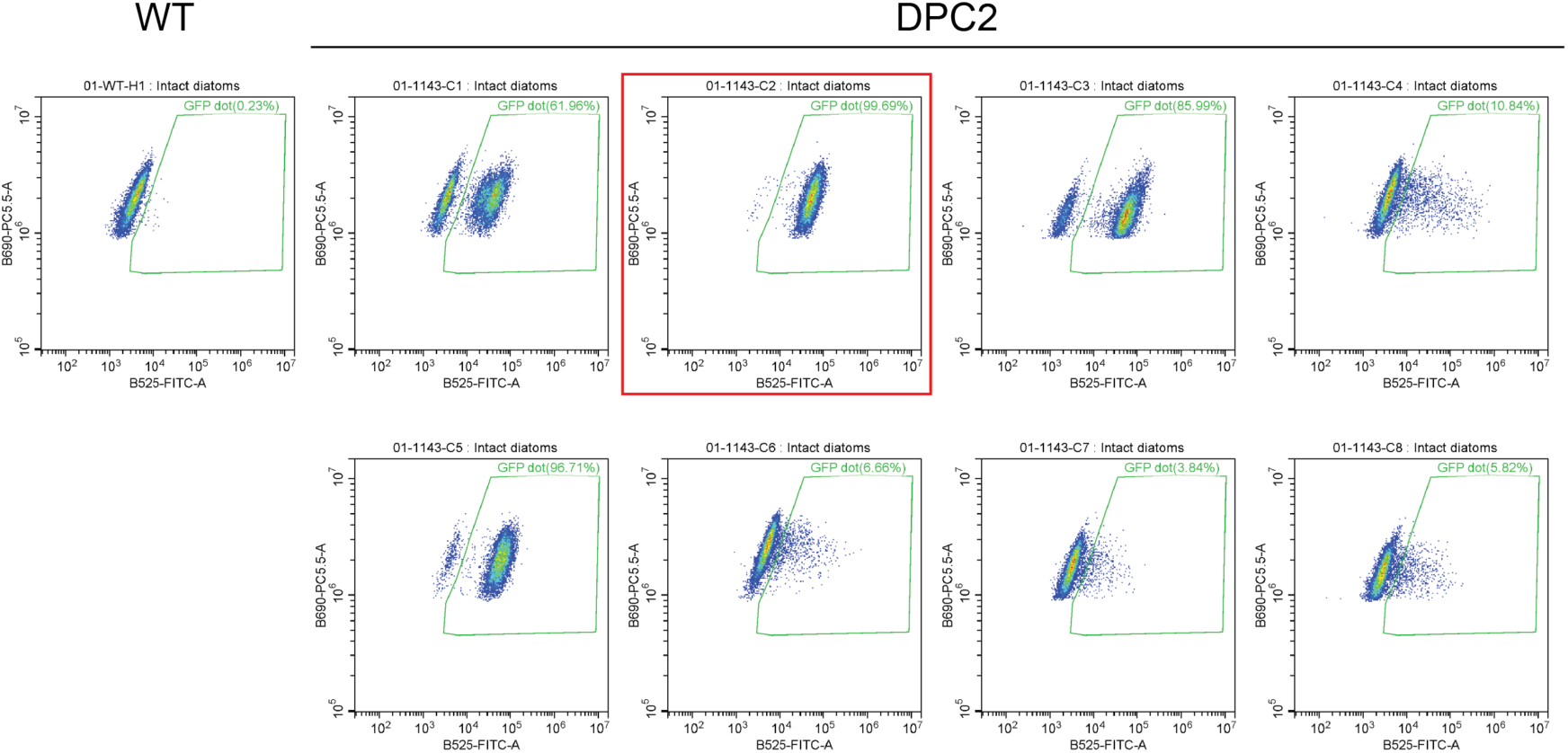
Flow cytometry screening to identify fluorophore fusion lines. Example screening data for Diatom Pyrenoid Component 2 (DPC2). Eight independent lines expressing DPC2-mEGFP from an episome were screened via flow cytometry. The green box outlines the gate used to quantify the percentage of cells that show mEGFP fluorescence with the percentage indicated above. The red box denotes the cell line used for subsequent imaging and APMS.

**Figure S3.**
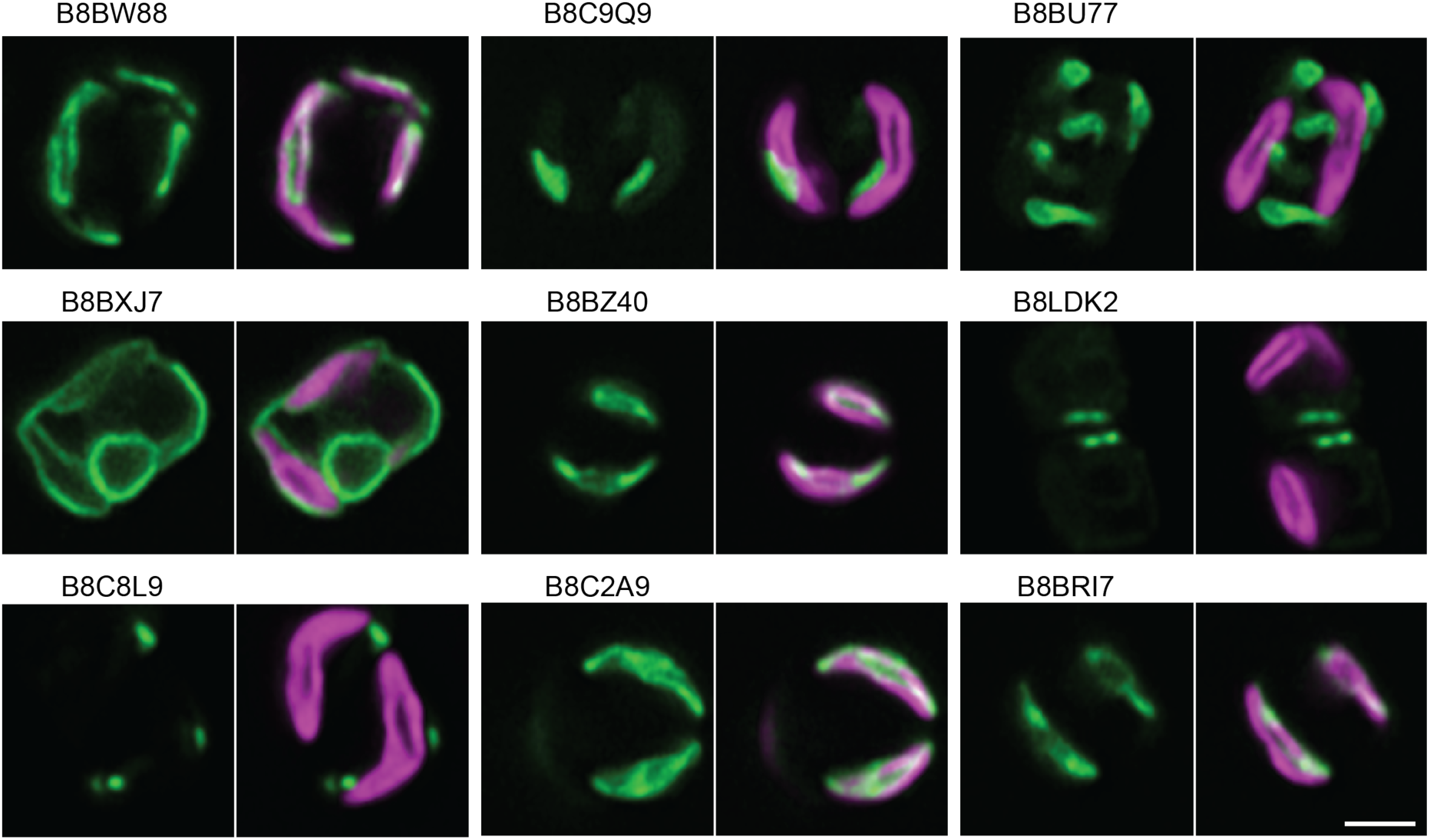
mEGFP fusion proteins that did not localize to the pyrenoid. Green: mEGFP fusion protein; Magenta: chlorophyll. Scale bar: 2 µm. Predicted functions/annotations: B8BW88, Aspartate/glutamate/uridylate kinase domain-containing protein; B8BXJ7, Uncharacterized protein; B8C8L9, Gfo/Idh/MocA-like oxidoreductase N-terminal domain-containing protein; B8C9Q9, Oxoglutarate/malate translocator; B8BZ40, Phosphoribulokinase; B8C2A9, S-malonyltransferase; B8BU77, Nucleoside-diphosphate kinase; B8LDK2, Glycoside hydrolase family 5 domain-containing protein; B8BRI7, 2-isopropylmalate synthase.

**Figure S4.**
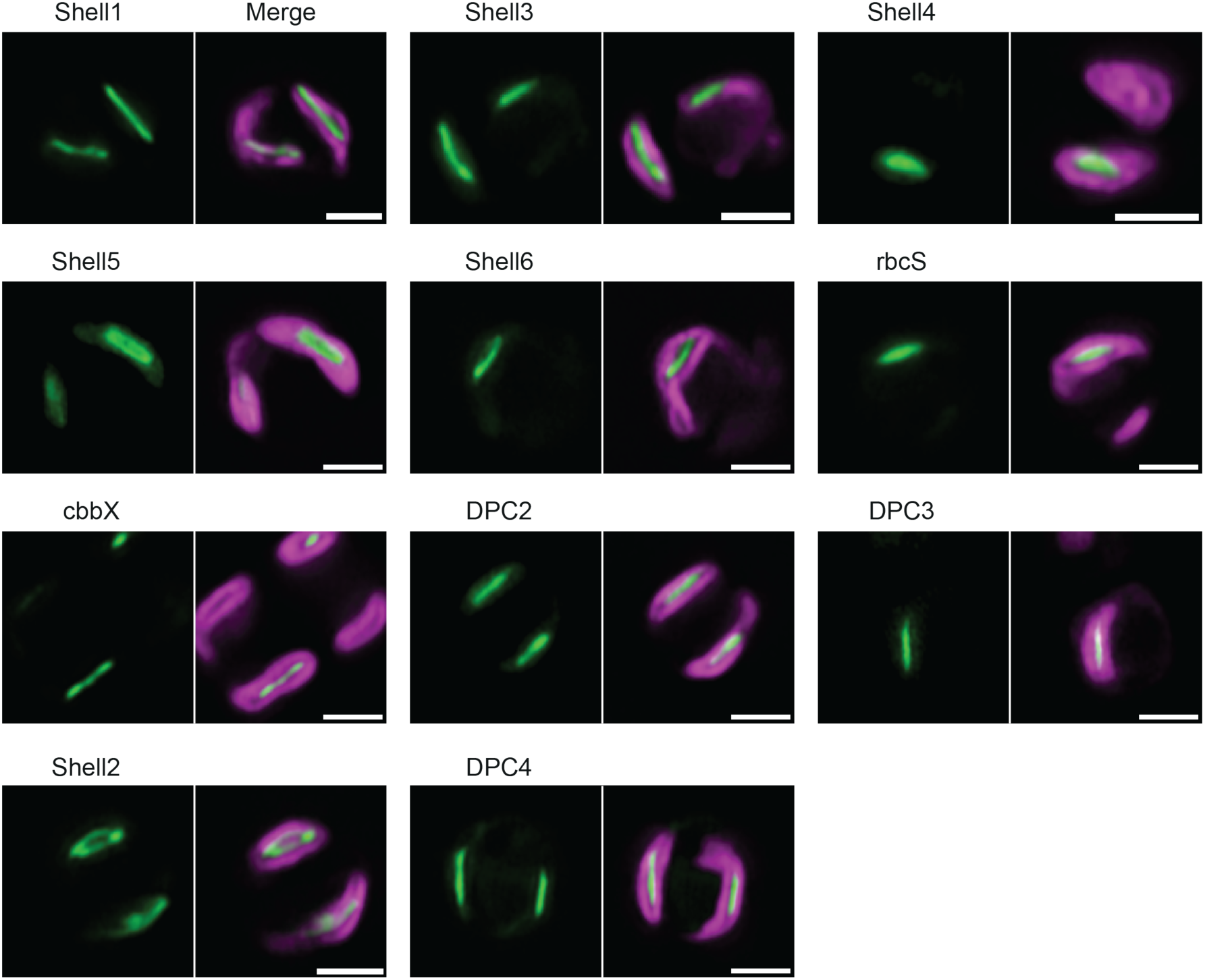
Images of additional mEGFP tagged lines. Images of additional pyrenoid localized proteins, and additional independently generated lines to those used in Fig. 2 and Fig. 3. Green: mEGFP fusion protein; Magenta: chlorophyll. Scale bars: 2 µm.

**Figure S5.**
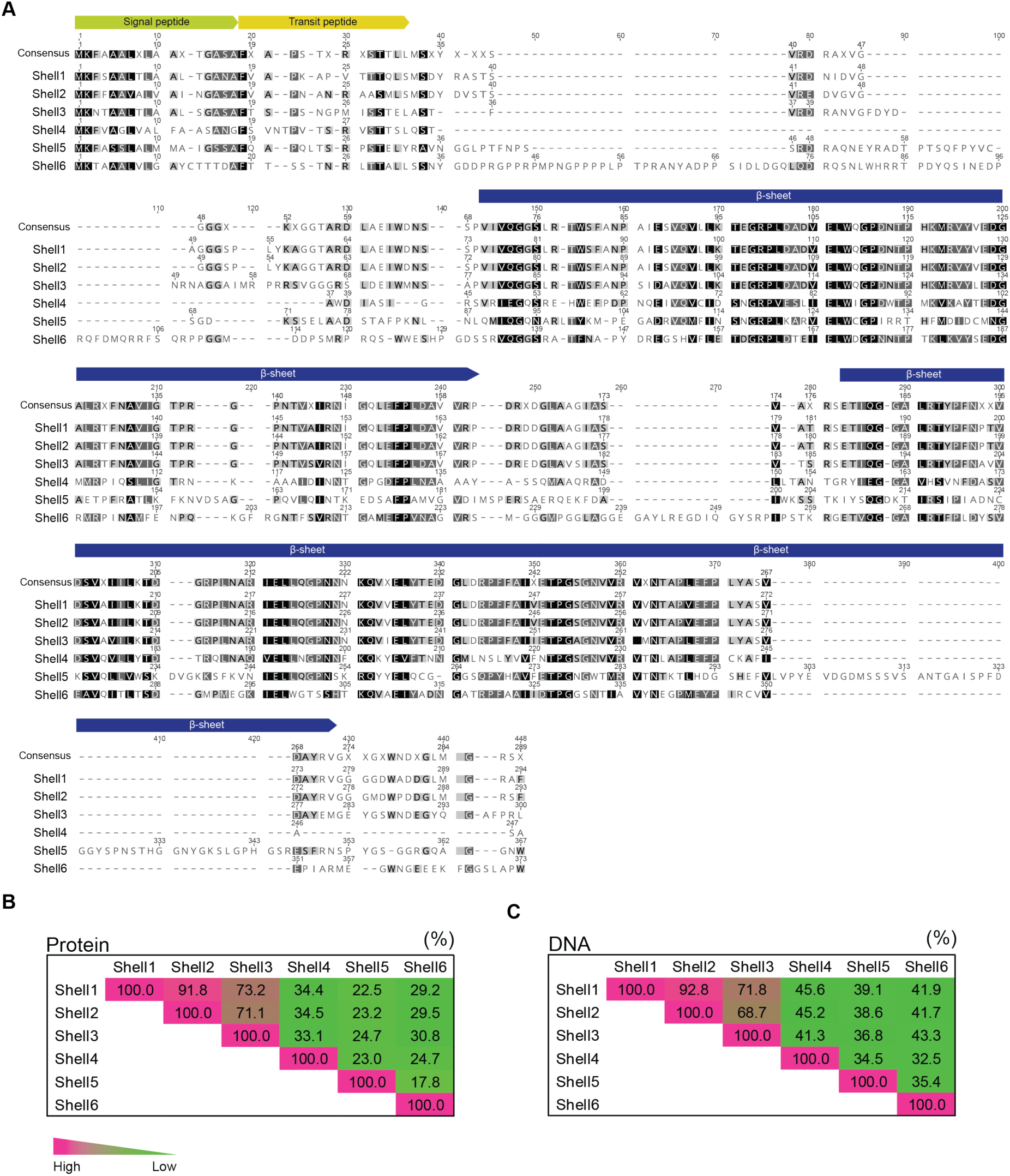
Shell protein amino acid alignment and sequence similarities. A. Alignment of full-length Shell amino acid sequences. B. Protein sequence similarities between Shell homologs, including predicted signal and transit peptides. C. Gene DNA sequence similarities between Shell homologs.

**Figure S6.**
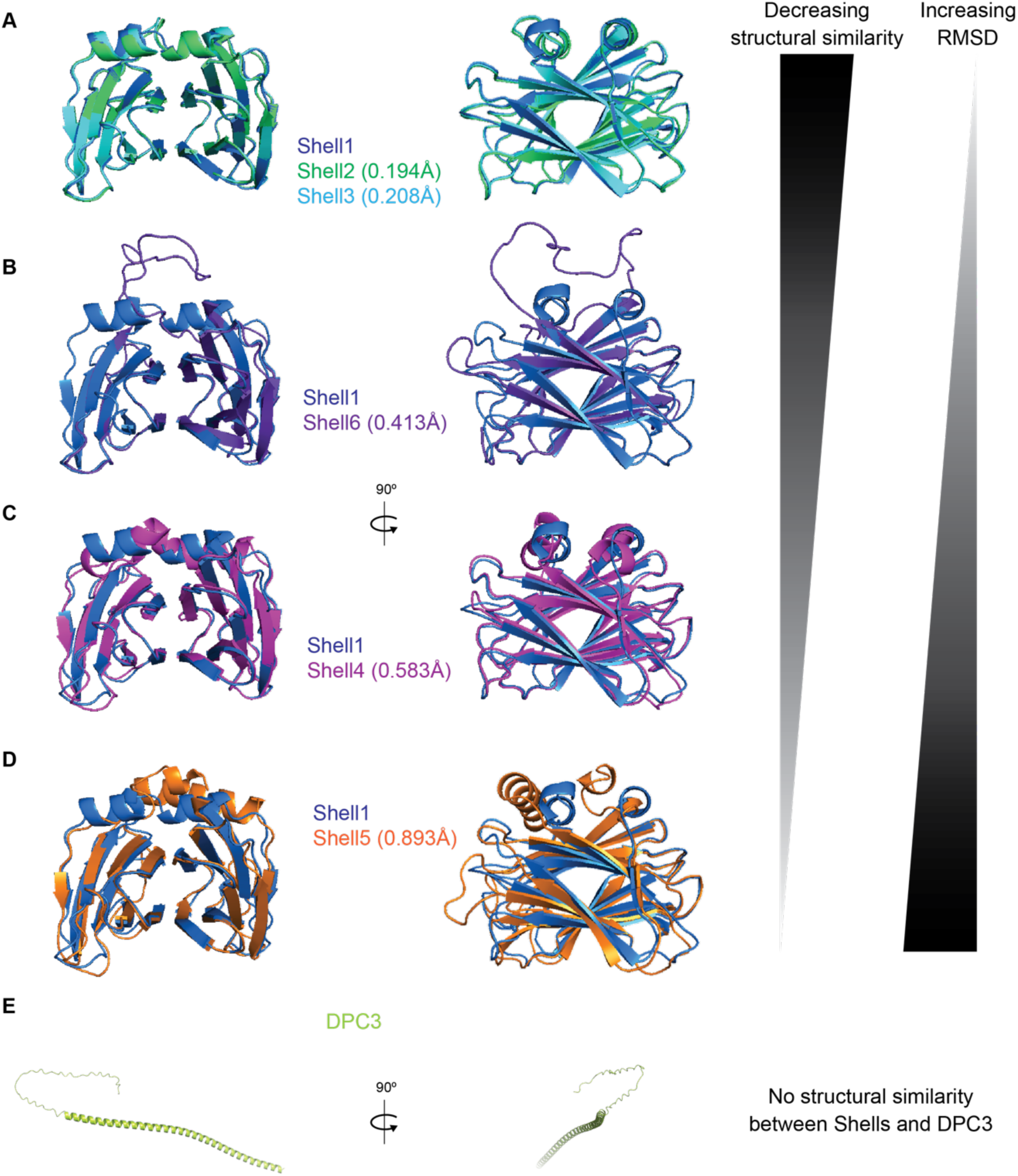
AlphaFold2 model comparisons of Shell proteins and DPC3. Root mean square deviation (RMSD) values relative to Shell1 are shown.

**Figure S7.**
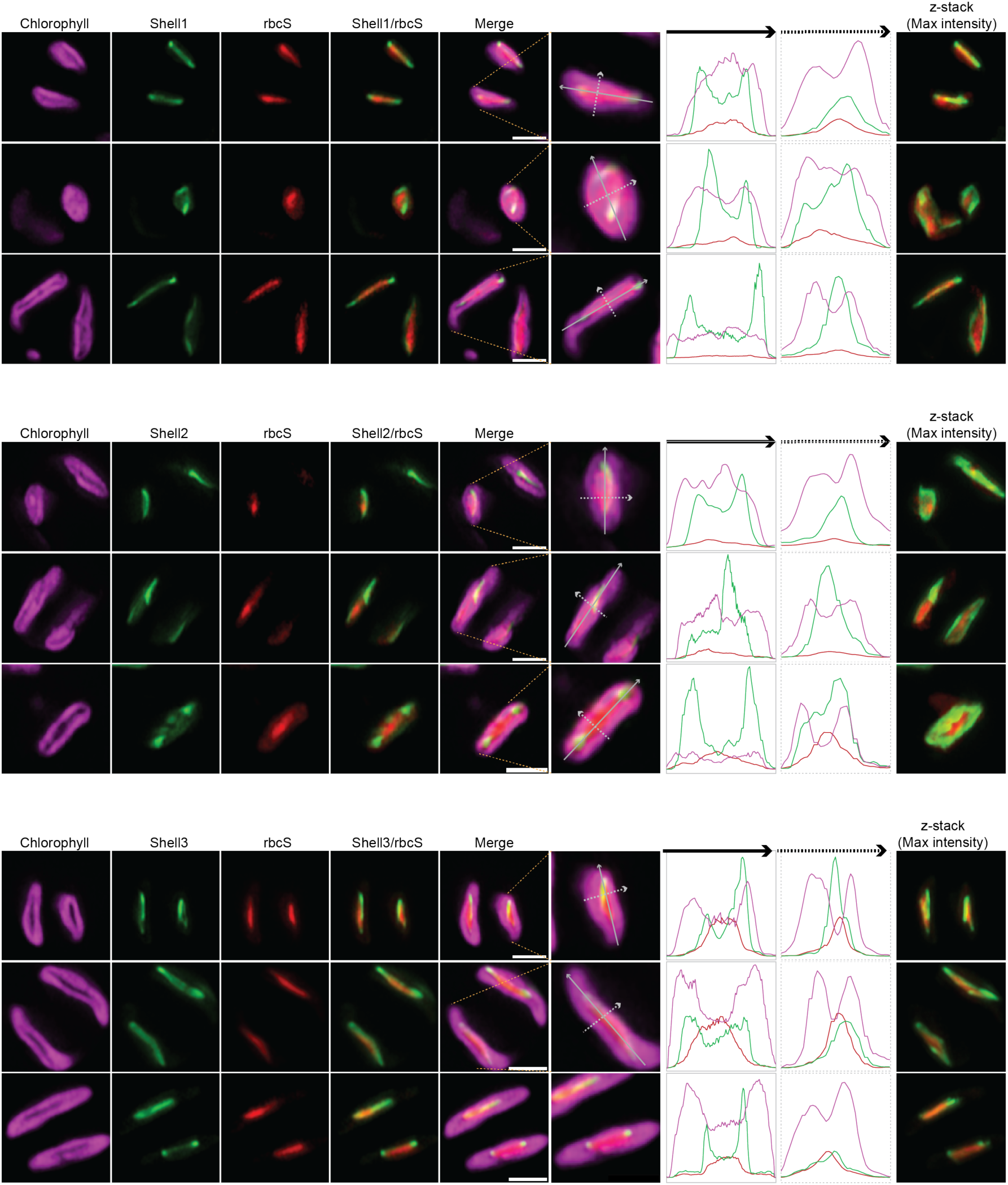
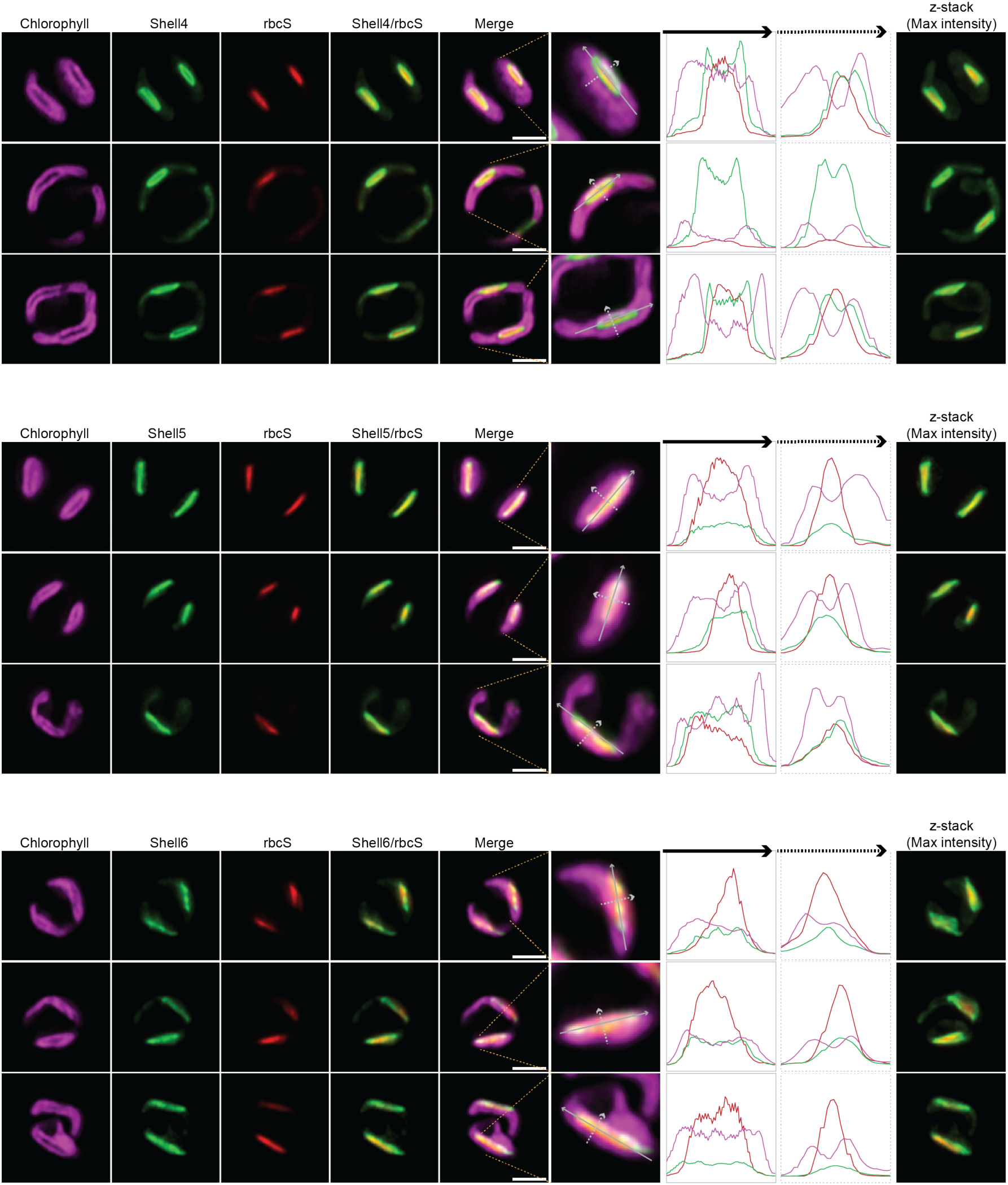
Co-localization of Shell proteins with Rubisco. Image columns 1-4: additional images of co-localization of Shell proteins with Rubisco. Scale bars: 2µm. Image columns 5-8: Fluorescence intensity cross-sections of chlorophyll, Shell and rbcS fluorescence across the pyrenoid. Level of Zoom changes between images. Image column 9: Z-stack max intensity projections.

**Figure S8.**
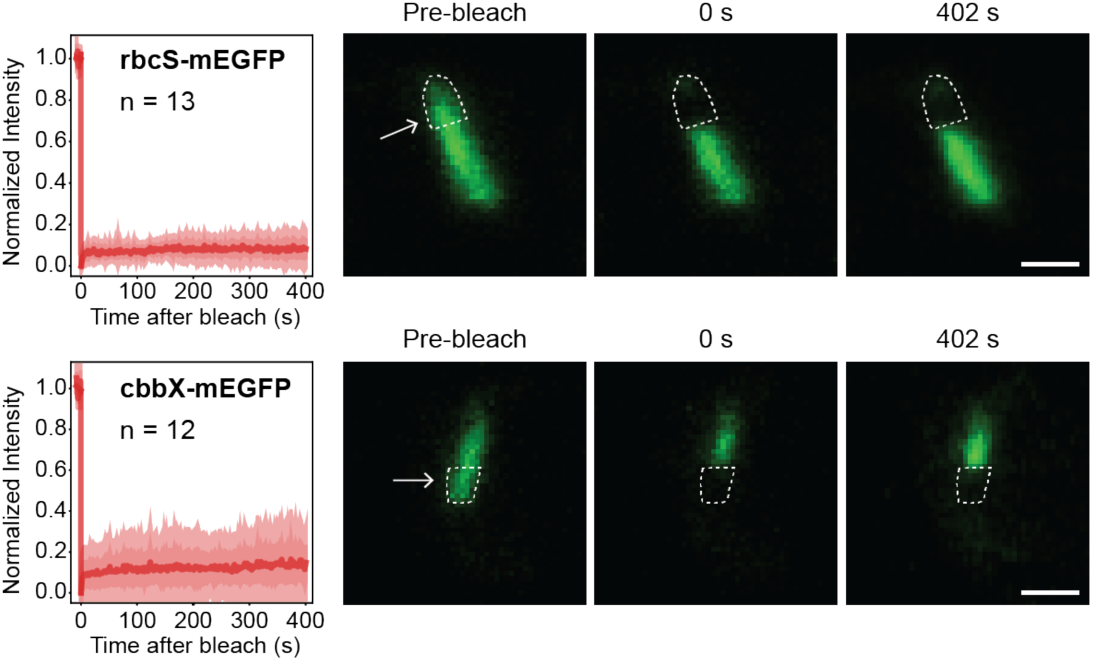
RbcS and cbbX are not mobile in the *T. pseudonana* pyrenoid. Fluorescence recovery after photobleaching (FRAP) experiments for rbcS and cbbX. Arrows indicate photobleached regions. Scale bars: 1 µm. Plots show S.E.M. and S.D. of the mean.

**Figure S9.**
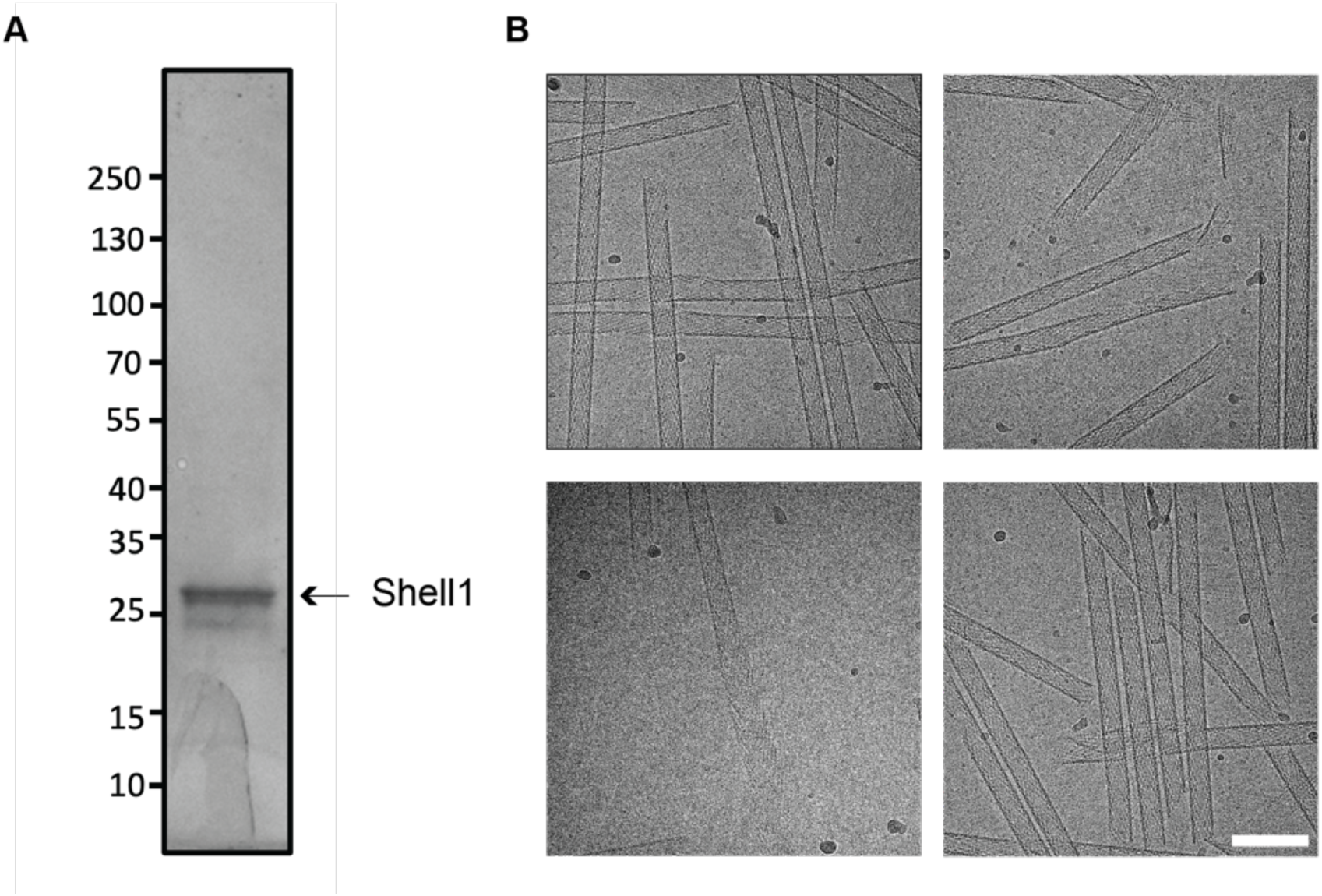
Shell1 forms tubes and sheets *in vitro*. A. Coomassie stained SDS-PAGE gel of recombinant Shell1 used for cryo-EM. B. Representative cryo-EM micrographs of Shell1 self-assembly into tubes and sheets. Scale bar: 100 nm.

**Figure S10.**
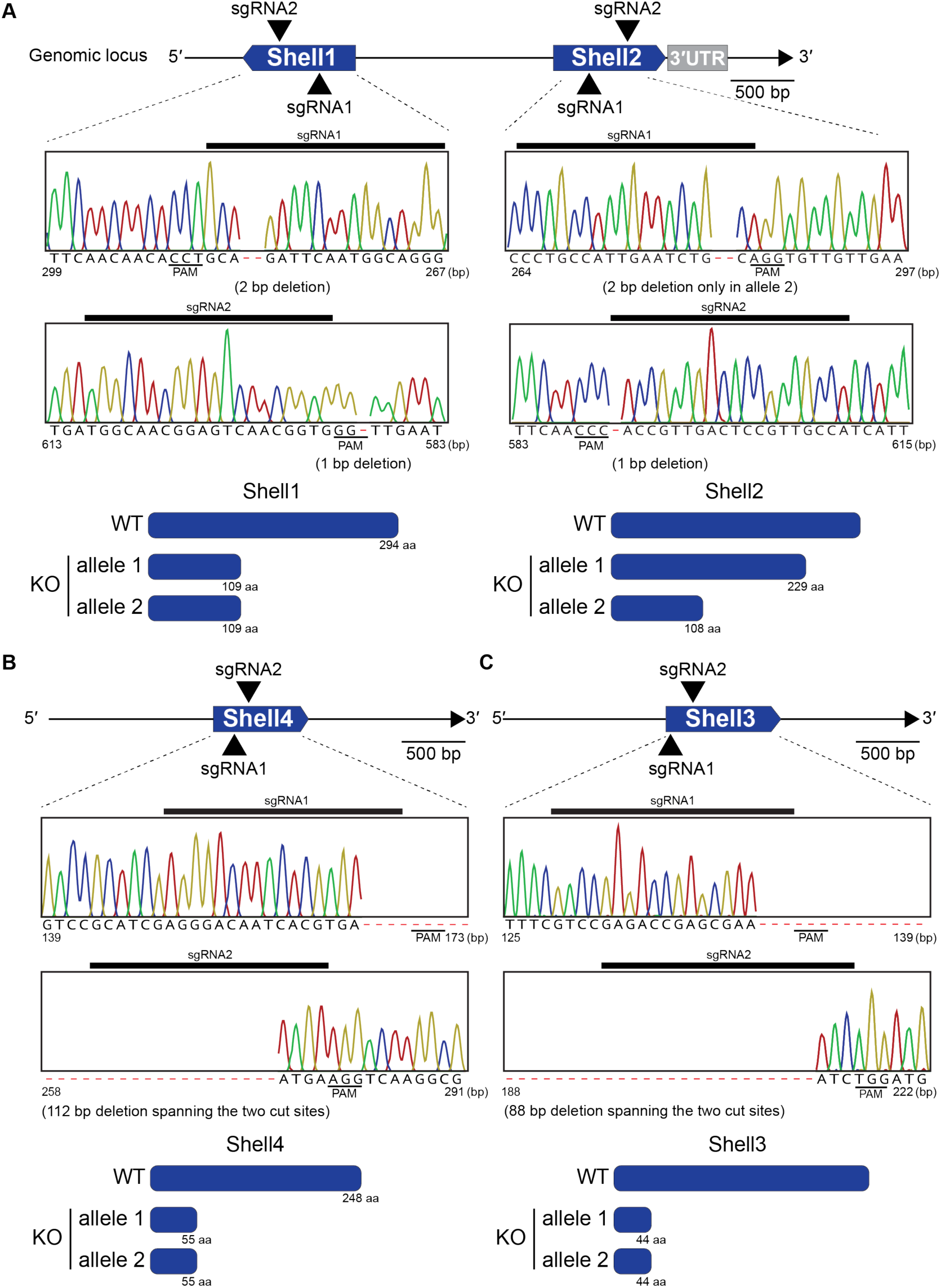
Shell1/2, Shell3, and Shell4 knock out genotyping. A. Due to 93% DNA sequence similarity between *Shell1* and *Shell2*, both genes were knocked out simultaneously by designing two sgRNAs that targeted conserved sequences in *Shell1* and *Shell2*. B. Biallelic knock out genotyping of Shell4. C. Biallelic knock out genotyping of Shell3.

**Figure S11.**
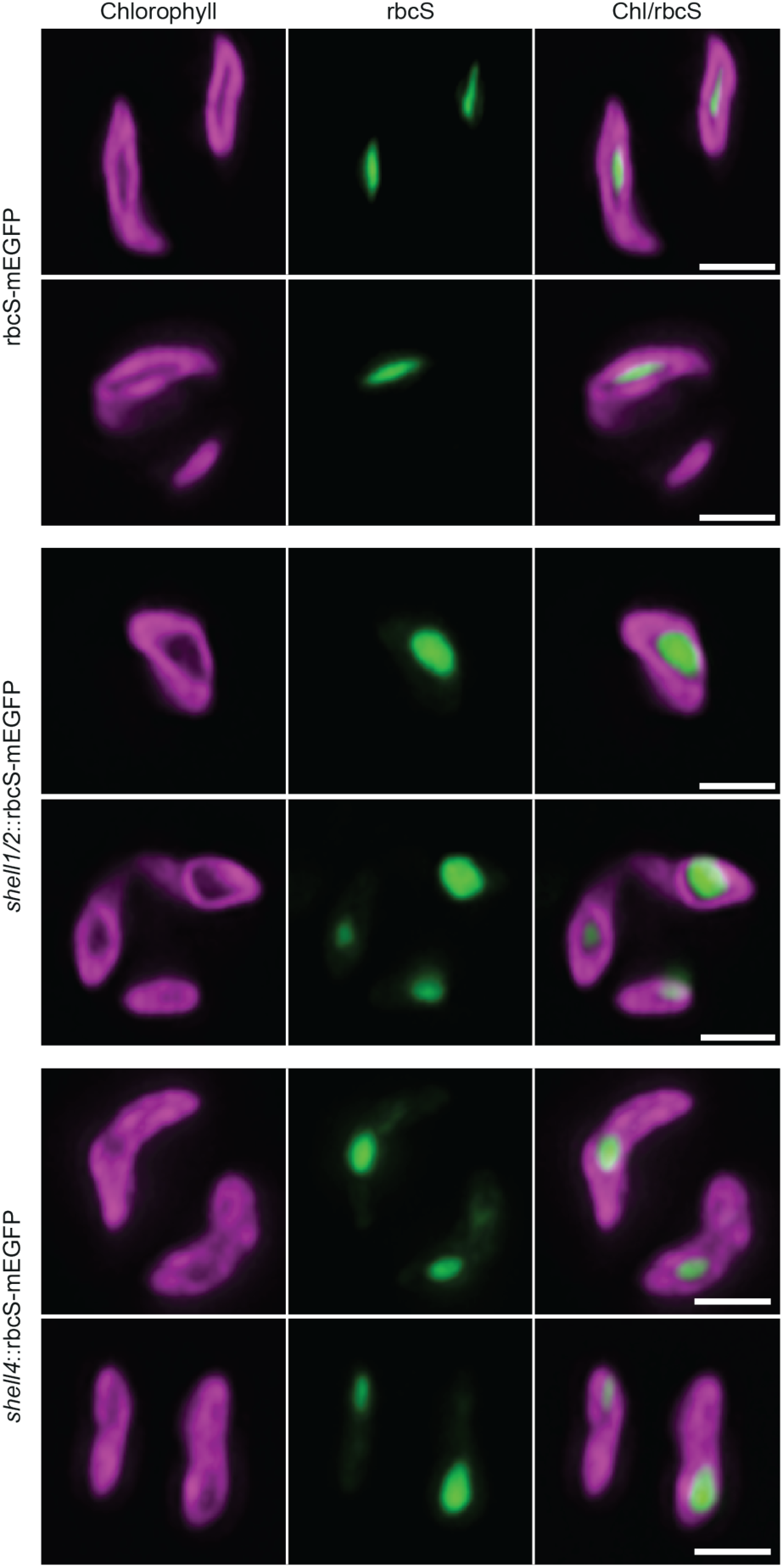
*Shell1/2* and *shell4* mutants have malformed pyrenoids. Additional images of WT::rbcS-mEGFP, *shell1/2*::rbcS-mEGFP and *shell4*::rbcS-mEGFP. Green: mEGFP fusion protein; Magenta: chlorophyll. Scale bars: 2 µm.

**Figure S12.**
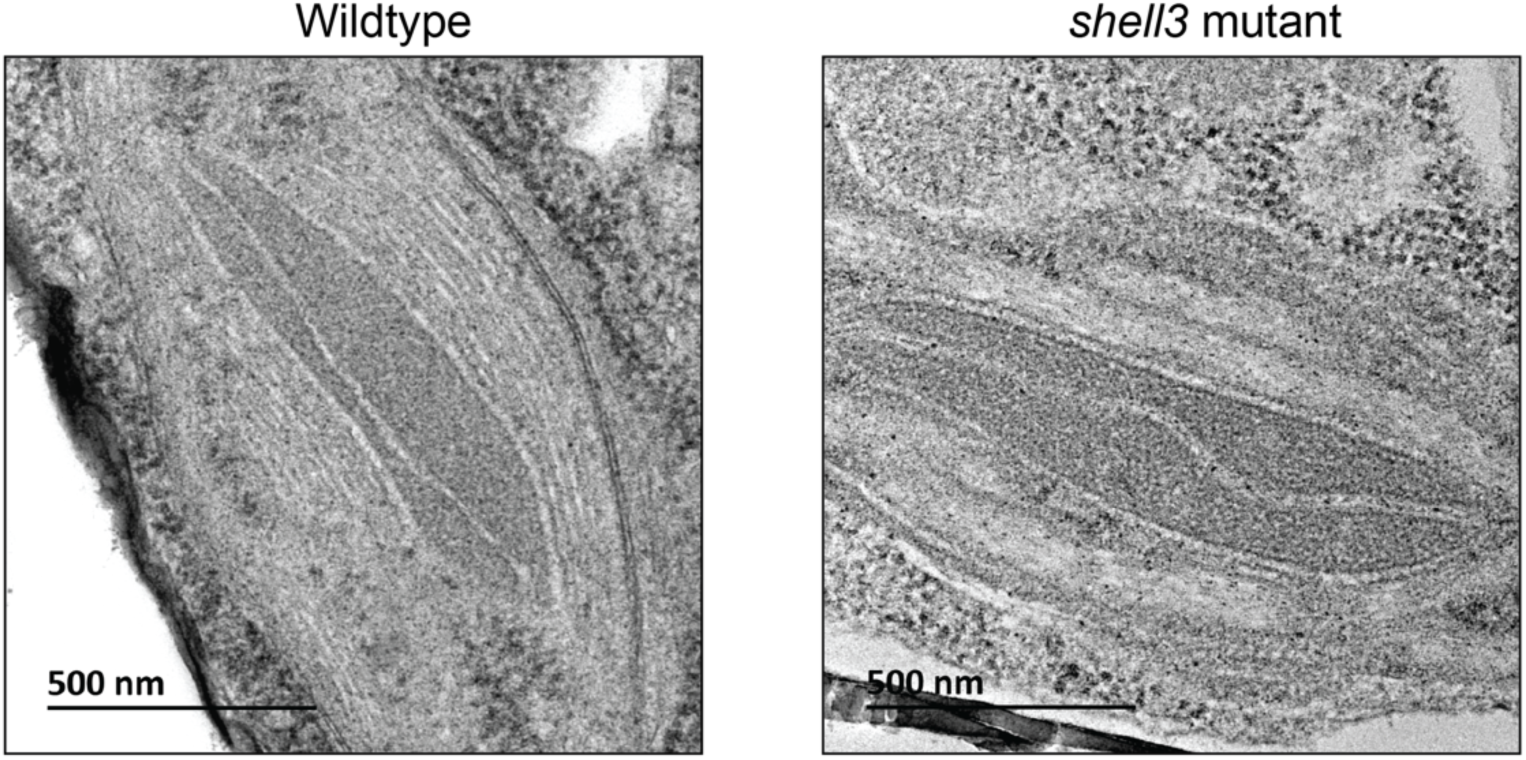
The *shell3* mutant has WT pyrenoid morphology. TEM images of WT and *shell3* mutant pyrenoids. The WT image is the same as used in Fig. 1.

**Figure S13.**
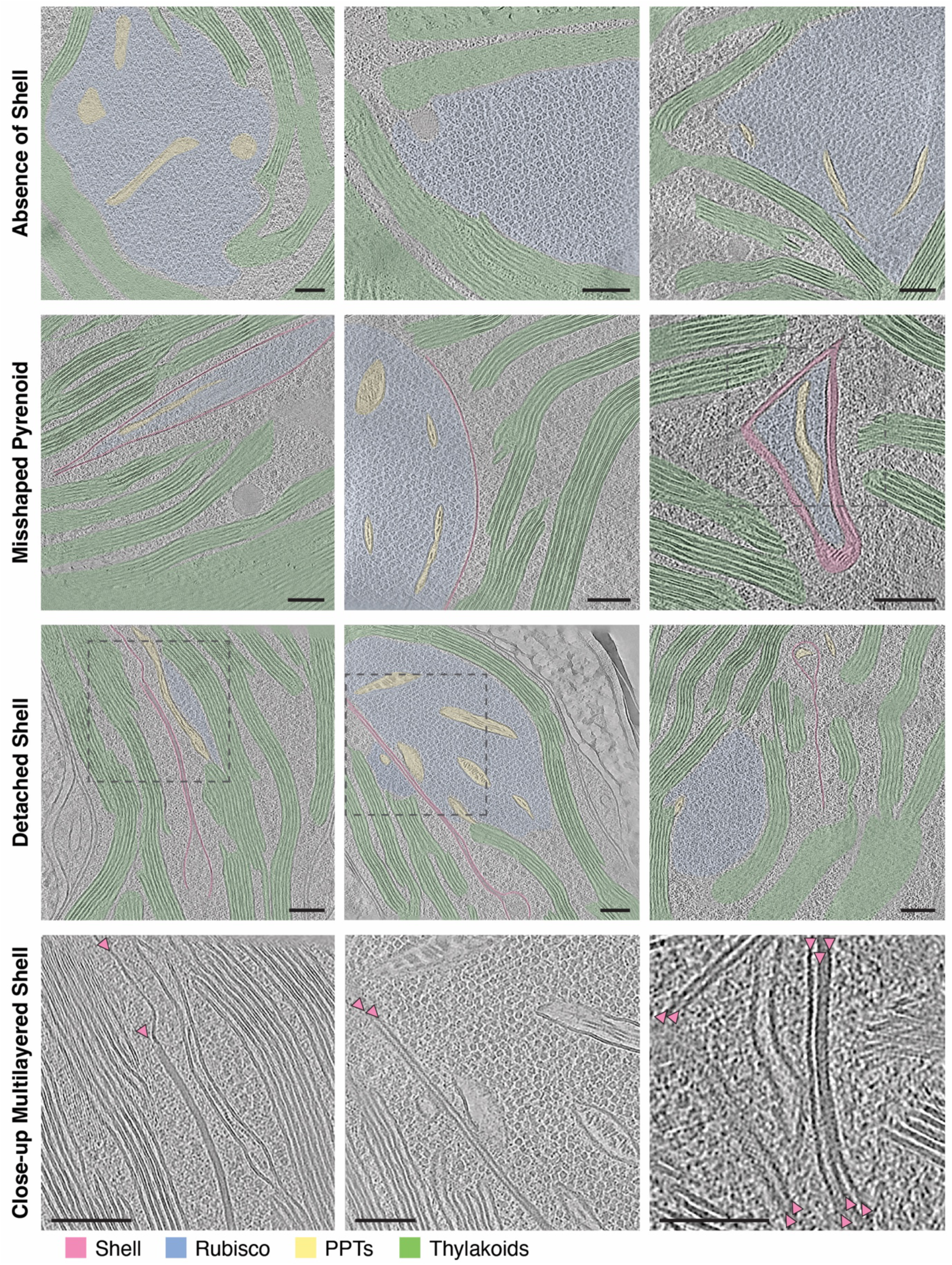
Additional tomograms of the *shell4* mutant pyrenoid morphologies. Additional tomogram slices of the different types of misassembled pyrenoids in the *shell4* mutant. The Shell (pink), Rubisco matrix (blue), thylakoids (green) and PPTs (yellow) are highlighted. Bottom row shows enlarge views of the boxed regions above. Scale bar: 100 nm

**Figure S14.**
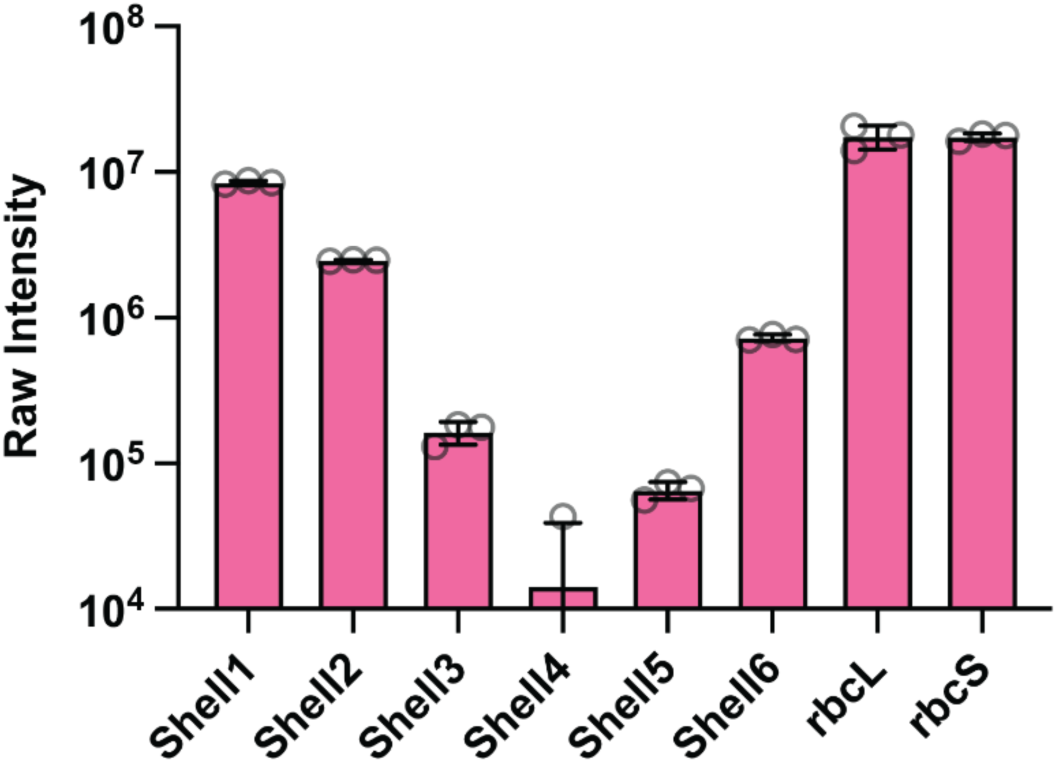
Whole-cell proteomics of the *shell4* mutant. Relative abundance of Shell proteins and Rubisco in the *shell4* mutant. Shell4 was detected at low levels in one of the three replicas. Error bars: S.D. of the mean.

